# Tandem duplications lead to novel expression patterns through exon shuffling in *D. yakuba*

**DOI:** 10.1101/097501

**Authors:** Rebekah L. Rogers, Ling Shao, Kevin R. Thornton

## Abstract

One common hypothesis to explain the impacts of tandem duplications is that whole gene duplications commonly produce additive changes in gene expression due to copy number changes. Here, we use genome wide RNA-seq data from a population sample of *Drosophila yakuba* to test this ‘gene dosage’ hypothesis. We observe little evidence of expression changes in response to whole transcript duplication capturing 5ʹ and 3ʹ UTRs. Among whole gene duplications, we observe evidence that dosage sharing across copies is likely to be common. The lack of expression changes after whole gene duplication suggests that the majority of genes are subject to tight regulatory control and therefore not sensitive to changes in gene copy number. Rather, we observe changes in expression level due to both shuffling of regulatory elements and the creation of chimeric structures via tandem duplication. Additionally, we observe 30 *de novo* gene structures arising from tandem duplications, 23 of which form with expression in the testes. Thus, the value of tandem duplications is likely to be more intricate than simple changes in gene dosage. The common regulatory effects from chimeric gene formation after tandem duplication may explain their contribution to genome evolution.

**Author Summary:** The enclosed work shows that whole gene duplications rarely affect gene expression, in contrast to widely held views that the adaptive value of duplicate genes is related to additive changes in gene expression due to gene copy number. We further explain how tandem duplications that create shuffled gene structures can force upregulation of gene sequences, *de novo* gene creation, and multifold changes in transcript levels.

These results show that tandem duplications can produce new genes that are a source of immediate novelty associated with more extreme expression changes than previously suggested by theory. Further, these gene expression changes are a potential source of both beneficial and pathogenic mutations, immediately relevant to clinical and medical genetics in humans and other metazoans.

## Introduction

Tandem duplications are known as a source of genetic novelty that can contribute new genes with novel functions (1, 2). However, after duplication, copies require many generations to facilitate functional divergence. The expected long wait times to develop new functions raise the risk that duplicate genes may be eliminated via non-functionalizing mutations before they can evolve new functions, even in large populations where effects of drift are limited (3). Indeed, loss appears to be the prevailing fate of duplicate and chimeric genes (4, 3, 5). One solution proposed for how duplicate genes might accumulate in genomes given these limitations is the duplication-degeneration-complimentation model (3). If duplicate genes accumulated very few mutations in regulatory sequences, they might partition expression profiles of duplicate copies. This expression divergence might drive a situation where neither copy could be eliminated, resulting in long term preservation in the genome (3). Similar models might also explain neofunctionalization as well (6). An alternative hypothesis to explain the utility of newly formed duplicates proposed that newly formed duplicate genes may contribute to expression variation through additive changes in gene expression due to gene dosage (7). More recently it has become possible to survey natural variation in gene expression at duplicated loci, in order to better distinguish the factors that contribute to the utility and maintenance of duplicate genes in the genome.

It is also less well understood how other types of constructs beyond whole gene duplications may contribute to regulatory and protein sequence diversity in nature. Chimeric genes and novel recruited UTRs can cause expression changes in novel tissues through the shuffling of regulatory elements (8, 9, 10, 11). Yet, previous surveys have simply looked at presence and absence of transcripts in tissues with no systematic survey of quantitative changes or have focused on small numbers of candidate genes. Similarly studies of CNVs in *D. melanogaster* have identified a role in eQTLs (12), but with assays in whole adult flies that do not resolve different types of regulatory changes or the precise mechanisms of such changes. Systematic, genome wide surveys of the effects tandem duplications produce on gene expression is essential as a first step toward understanding how duplicate genes may contribute to regulatory variation in natural populations. *D. yakuba* offers an excellent genetic model to examine changes in genome architecture and genome content in natural populations. Comparisons across the *Drosophila* genus indicate that *D. yakuba* has experienced a large number of changes in genome structure (13), and population level surveys have identified large numbers of duplications that are polymorphic in comparison with sister species (14).

Here, we describe a genome wide survey of polymorphic variation for tandem duplications in natural populations of *D. yakuba* and the types of regulatory changes that they can facilitate. We further describe biases in the ancestral expression patterns of genes that are duplicated. We show that whole gene duplications rarely produce effects on expression. In order to survey the detailed changes in gene expression produced by chimeric genes, gene fragments and recruited non-coding sequence, we introduce a hidden Markov model to assay site specific changes in gene expression, independent from gene annotations. These mutations form new gene structures not reflected in reference genome annotations, requiring an alternative approach from existing differential expression testing software. Using this new model, we identify 30 cases where duplications result in *de novo* gene origination, with an excess of new genes appearing with expression in the testes. Tandem duplications associated with chimeric constructs, novel UTRs, and recruited non-coding sequence are commonly associated with regulatory changes. These findings are consistent with previous studies showing testes bias (15). The results presented here suggest that complex changes in gene structures will be an important source of mutations of major effect and that the value of whole gene duplications is unlikely to lie in additive changes in transcript levels due to gene copy number.

## Results

Many newly formed tandem duplicates are associated with non-neutral effects (16, 17, 18, 19, 20,21, 16), in contrast with theoretical claims that tandem duplications are likely to be nearly neutral (3, 1). Yet, the reasons behind these non-neutral impacts are unclear. Here, we describe expression data for tandem duplications as a first step to elucidate the extent to which the molecular impacts of tandem duplications may explain their functional and evolutionary impacts. Using high coverage genomic sequence data we previously identified tandem duplications in population genomic samples for *D. yakuba*, with high validation rates of 97%, for duplications ranging from 74 bp to 25,000 bp in length (14). We performed RNA-sequencing for adult male and female soma and reproductive tissues in 15 sample strains of *D. yakuba* as well as three replicates of the *D. yakuba* reference, which contains none of these tandem duplications. We have assayed transcript levels in new RNA-seq data for 15 of the 20 sample strains from Rogers et al, 2014 (14) as well as previously published data for 3 replicates of the reference strain (20) to obtain a portrait of regulatory changes that complex mutations can produce. Among strains assayed with RNA-seq data, we have identified 1116 tandem duplications in total. Among the 1116 duplications, 112 capture solely intergenic sequence while 1004 tandem duplications capture a total of 1306 genes or gene fragments based on new RNA-seq based gene annotations (22). Among these, we identify 66 whole gene duplications, 76 chimeric genes, and 30 cases of recruited non-coding sequences that might potentially contribute to *de novo* gene formation.

### Scarce support for the Dosage Hypothesis

One commonly proposed source of adaptive variation suggests tandem duplications may cause two-fold changes in transcript levels, resulting in quantitative phenotypic change via “gene dosage” (23, 12, 24, 7). This “dosage” hypothesis offers one putative genetic mechanism for immediate evolutionary change prior to pseudogenization and loss. However, we observe scarce support for changes in RNA levels within tissues in response to duplication using both quantile normalized expression data (Figure 1, Figure S1) and FPKM normalized expression data (*P* ≥ 0.37; Figure S2). Using the Tophat/Cufflinks differential expression testing suite, we assayed 52 whole gene duplications (including UTRs) that had gene models that passed cuffdiff quality filters. In every tissue, the number of genes with significantly increased expression levels compared to the reference strain was not significantly different from genome wide expectations (Table S1). In all of these cases, expression levels did not reflect additive two-fold changes in expression levels but rather indicated much greater fold change (Figure S3, Table S2). When we require at least 1 kb of upstream and downstream sequence, we do not observe any evidence of additive changes in gene expression. This is equally true when restricting duplications to cases where reference expression level is FPKM≥ 2. Cufflinks is fully capable of detecting low level changes in gene expression (25). The whole gene duplications with upregulated expression here are associated with several different functions with no clear functional enrichment. Variants include testes expressed endopetidases, a metalloendopeptidase, a chorion protein, and two metabolism genes: sorbitol dehydrogenase, giberellin oxidase (Table S3). However it is not clear that any of these expression changes are the product of duplication. High frequency duplications may be older and have secondary modifications on expression levels. They may also be filtered by selective pressures in comparison with low frequency duplications, possibly weeding out genes with expression changes. We examined 33 singleton variants that are expected to reflect primarily newly formed duplications, including detrimental (but not lethal) variants. Qualitatively, results remained unchanged, with no significant excess of expression changes for whole gene duplications (Table S4). Thus, there appears to be little support for this gene dosage hypothesis for duplicate genes in adult tissues.

**Figure 1:**
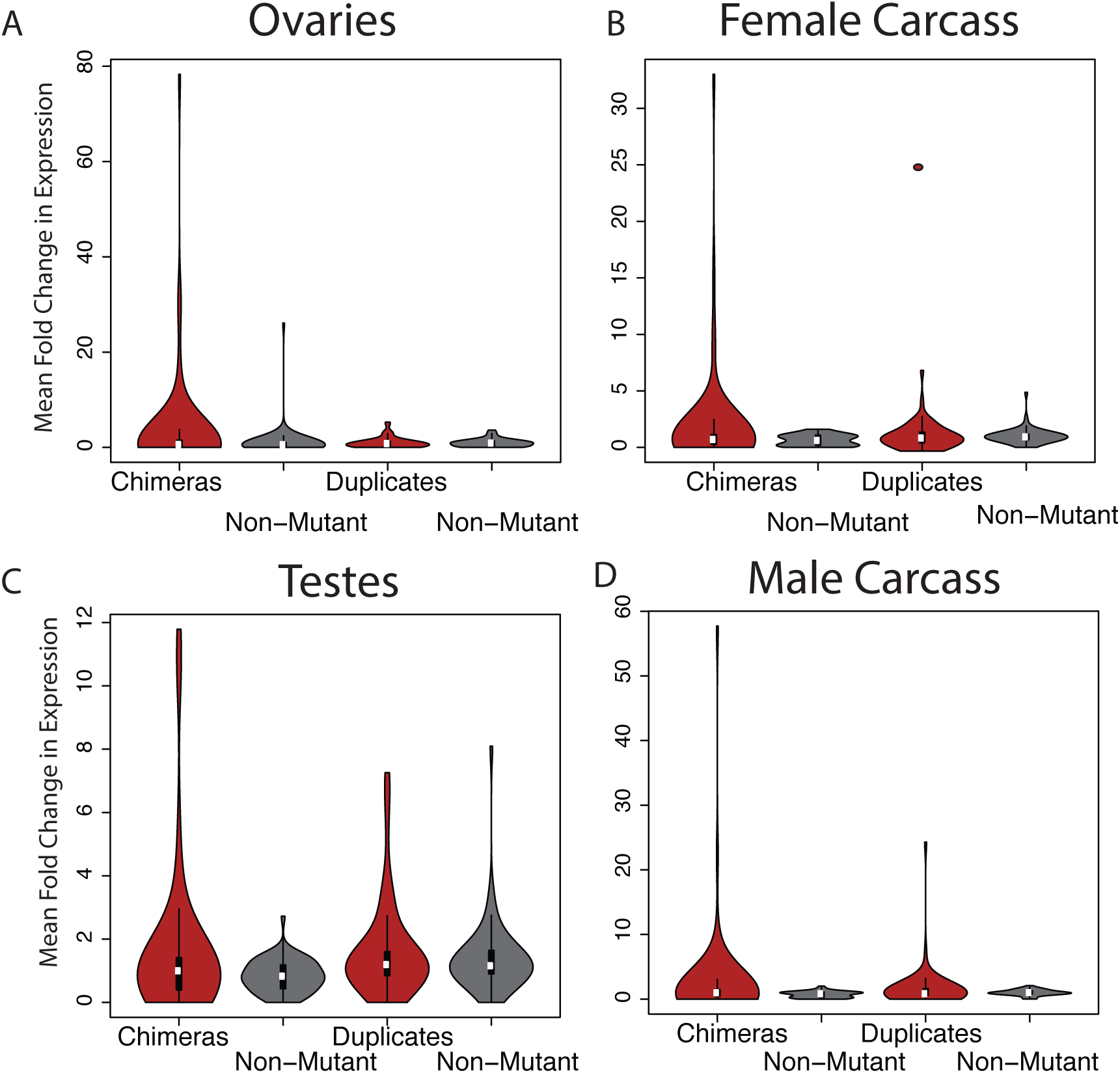
Mean fold change for chimeric genes in sample strains vs. reference for strains containing chimeras or whole gene duplicates (red) and unmutated sample strains for the same regions (grey). Chimeric genes are more likely to result in high mean fold change than unmutated counterparts in all tissues. Whole gene duplicates create multifold expression changes more rarely.

One hypothesis for the lack of increased expression is that secondary silencing of additional copies might subdue expression changes produced by whole gene duplication. We identified 52 whole gene duplications with at least one ‘heterozygous’ SNP mutation present that might differentiate duplicate copies based on genomic sequencing. We filtered out SNPs that display asymmetric expression in non-duplicate strains, which would indicate allele-specific expression independent of duplication. This leaves a remaining 11 candidates that might represent asymmetric expression of duplicate genes in at least one tissue (Table S5-S6), though the possibility of allele specific expression at a single locus cannot be ruled out. These numbers represent a minority of whole gene duplications. Thus, we conclude that whole gene duplication with dosage-sharing is common.

Recent work has found some evidence for increases in expression at CNVs, in contradiction with the data presented here (26). It is possible that what they describe as complete duplications do not include UTR sequences, mis-identifying chimeric constructs, which we show are commonly associated with expression effects. It is also possible that their filters only for highly expressed genes focus on genes that are more likely to be limited by transcription. Finally, their permutation test controls for a gene-specific p-value of 0.05, but does not control for the genome-wide false positive rate. It is unclear which of these explanations may clarify the discrepancy between this dataset for *D. melanogaster* and the data presented here for *D. yakuba*.

### Gene expression changes from alternative gene structures

In light of these surprising results, we determined to take a closer look at the expression impacts of these tandem duplications, especially alternative gene structures beyond whole gene duplication. Chimeric gene structures, gene fragments, and cases of recruited non-coding sequence all reflect partial gene changes, not present in reference GFF files. Precise breakpoints for most tandem duplications cannot always be determined (14) even with high confirmation rates in PacBio long molecule data. To identify more detail with respect to changes in gene expression for alternative gene structures whose precise breakpoints remain unresolved, we developed a hidden Markov model to identify changes in gene expression for individual sites in the genome. This HMM allows for differential expression testing for segments of chimeric genes, gene fragments, and cases of recruited non-coding sequence. The method is agnostic with respect to size of genetic constructs assayed and it does not require perfect knowledge of duplication breakpoints, in contrast with standard differential expression testing software. To establish a baseline for comparison, we used the HMM to identify gene expression changes at whole gene duplications. In total, a maximum of 5 out of 66 whole gene duplications that capture both UTRs display signals of increased expression for 50% or more of total exonic sequence (Figure S3; Table 1) whereas the majority of genes remain unchanged (e.g. *GE18452*, Figure 2). Most promoters in *Drosophila* lie within 50 bp of gene sequences (27). Restricting whole gene duplications to cases where 100 bp of upstream and downstream of both UTRs where the promoter is likely to be captured, 5 out of 58 sequences display expression changes. Both with and without upstream regions the likelihood of upregulation is not significantly different from the background rate of 5.26% (SI Appendix, Table S7; 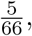 *P* = 0.7787; binomial test 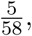 *P* = 0.2324). The HMM used to identify expression differences is fully capable of detecting 2x expression changes (SI Appendix, Figure S4), suggesting that the lack of genes with expression changes is not solely due to a lack of power. Both the number of whole gene duplications identified as upregulated and the background rates of upregulation are lower than results from cuffdiff, but both methods suggest that whole gene duplication is not associated with additive increases in expression where two copies of a gene produce a greater number of transcripts. Only one gene is identified as upregulated in male carcass, and this locus also exhibits upregulation in female carcass. Hence, it is unlikely that the use of paired end reads in male tissues has a strong influence to produce higher power in the HMM. No gene ontology functions are overrepresented among the five genes (Table S3).

**Figure 2:**
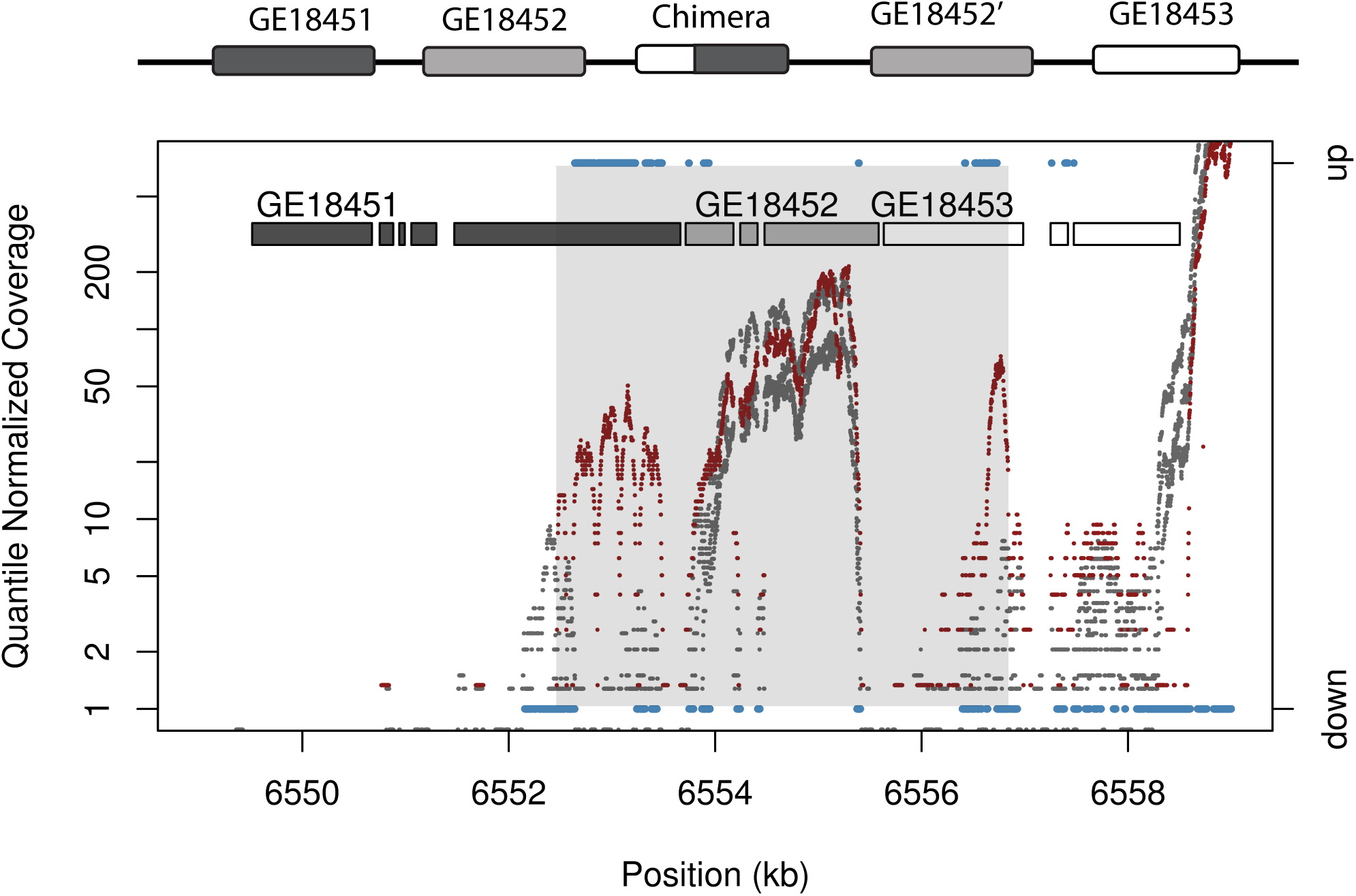
Chimeric gene structures result in novel expression patterns. A tandem duplication that does not respect gene boundaries unites the 5ʹ end of *GE18453* with the 3ʹ end of *GE18451* to produce a chimeric gene on chromosome 2L. Plot shows quantile normalized coverage in RNA seq data for sample (red) and reference (grey) with HMM output (blue) on chromosome 2L for female carcass. The chimera displays a change in transcript levels, while transcript levels for parental gene sequence are not altered. Sites with upregulated or downregulated sequence as defined by HMM output is shown in blue, using the right axis. HMM state calls for sites with unchanged expression are not shown. The region spanned by the tandem duplication is shaded in grey. The region spanned by the chimeric gene shows high-level upregulation. The whole gene duplication of *GE18452* does not display a significant change in mRNA levels but rather falls within the bounds of expression profiles for reference replicates (Ref FPKM=19.9; Sample FPKM=24.5; uncorrected *P* = 0.52; corrected *P* = 1.0).

**Table 1:** Upregulated genes

|  |  |  |  |
| --- | --- | --- | --- |
| Chimeras | Tissue | Upregulated | Total |
|  | Female Carcass | 5 | 76 |
|  | Female Ovary | 11 | 76 |
|  | Male Carcass | 10 | 76 |
|  | Male Testes | 7 | 76 |
|  | Aggregate | 24 | 76 |
| Whole Gene | Tissue | Upregulated | Total |
|  | Female Carcass | 3 | 66 |
|  | Female Ovary | 2 | 66 |
|  | Male Carcass | 1 | 66 |
|  | Male Testes | 0 | 66 |
|  | Aggregate | 5 | 66 |
| Whole Gene and 100 bp Intergenic | Tissue | Upregulated | Total |
|  | Female Carcass | 3 | 58 |
|  | Female Ovary | 2 | 58 |
|  | Male Carcass | 1 | 58 |
|  | Male Testes | 0 | 58 |
|  | Aggregate | 5 | 58 |
| *de novo* | Tissue | Upregulated | Total |
|  | Female Carcass | 7 | 1116 |
|  | Female Ovary | 2 | 1116 |
|  | Male Carcass | 10 | 1116 |
|  | Male Testes | 23 | 1116 |
|  | Aggregate | 30 | 1116 |

We observe one case where a duplication followed by a secondary deletion (Figure S5) (14), has resulted in upregulation of a gene fragment only at the modified locus, not the faithfully copied parental gene, showing that complex mutations can produce regulatory changes when RNA-level is unaltered at the unmodified paralog (Figure 3). Coverage from whole genome Illumina sequencing libraries of genomic DNA (14) shows a two-fold to three-fold increase in coverage for the portion of the duplicated segment not affected by the deletion, indicating that this segment is not multi-copy to a level that would explain the observed expression change (SI Appendix, Figure S5). Tandem duplications that do not respect gene boundaries can also create chimeric gene sequences via exon-shuffling (28) (SI Appendix, Figure S6A). In contrast to whole gene duplications, chimeric gene structures often result in expression changes. Among the 15 lines we identified 76 chimeric genes arising from tandem duplication. Of these a total of 24 chimeras display increased expression for 50% or more of exonic sequence within the duplicated gene segment (either 5ʹ or 3ʹ). These numbers are significantly different from random expectations given a background rate of 5.26% (binomial test 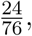 *P* = 5.16 × 10^−13^). The high mean fold change across all sites captured in chimera formation indicates high levels of upregulation independently from HMM results regardless of the tissue assayed (Figure 1).

**Figure 3:**
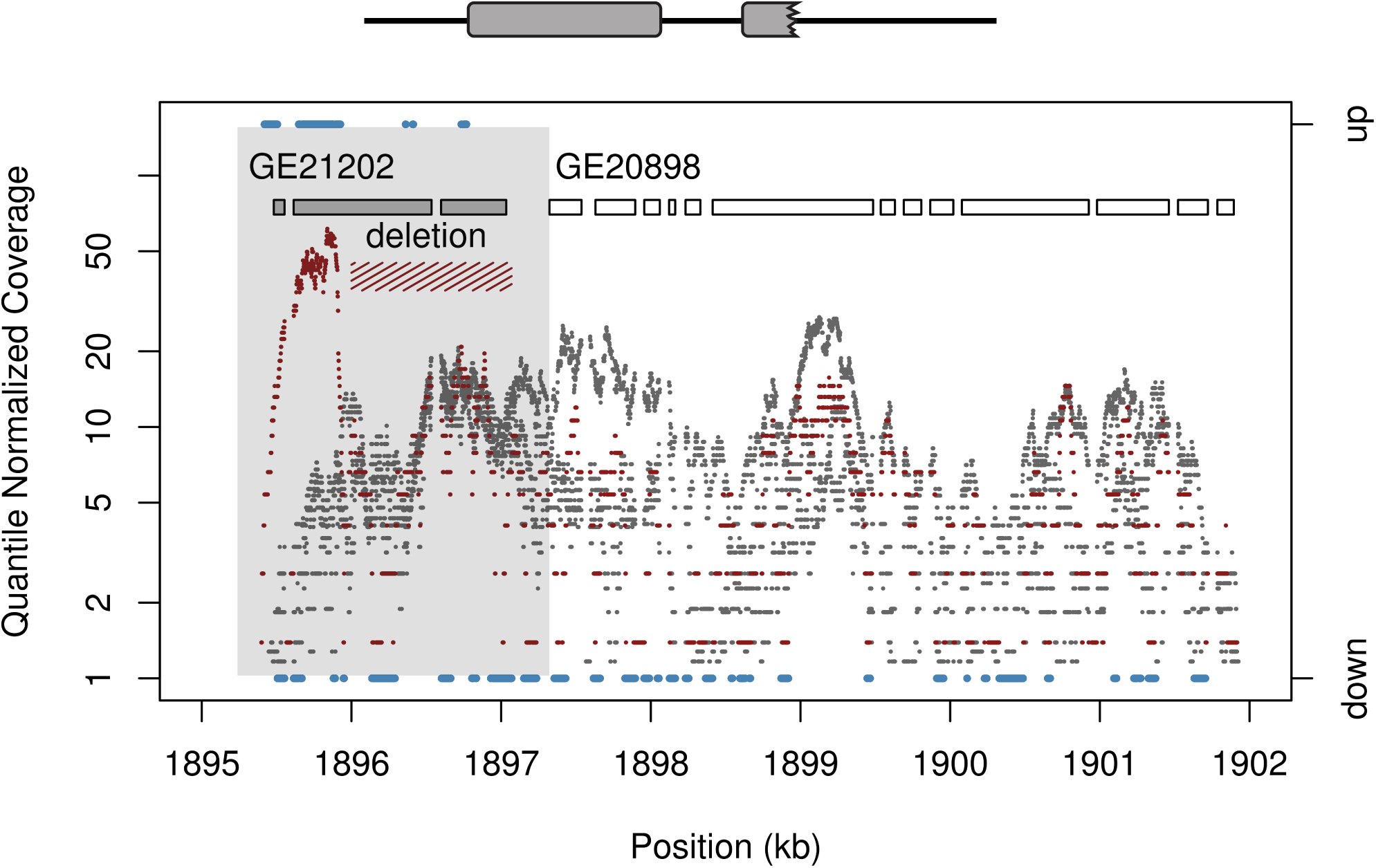
Duplication followed by secondary deletion, as indicated by a total of 104 long-spanning read pairs, leads to an expression change in a gene fragment of *GE21202* on chromosome 3L. Plot shows normalized coverage in RNA seq data for sample (red) and reference (grey) with HMM output (blue) on chromosome 3L. Only the sample strain with the deletion shows such upregulation. Transcript levels increase by greater than two-fold, beyond changes that would be produced by additive changes in gene dosage. Sites with upregulated or downregulated sequence as defined by HMM output is shown in blue, using the right axis. HMM state calls for sites with unchanged expression are not shown. HMM output for upregulated regions match well with the predicted gene structures formed by this complex mutation. The region spanned by the tandem duplication is shaded in grey.

These changes in gene expression are not consistent with additive effects of gene dosage, but rather reflect gene upregulation above two-fold changes due to the shuffling of regulatory elements in 5ʹ and 3ʹ segments of the gene. Plots of RNA-seq coverage and HMM output for these regions reflect the changes in gene structure, with only regions matching to chimeras exhibiting expression changes, not parental genes (Figure 2). These results suggest that expression changes are a direct product of chimera formation, not of environmental variation or secondary mutations that alter gene expression. Even with substantially less stringent criteria allowing for any expression change at least 50 bp in length, chimeric genes have a larger percentage of expression effects than whole gene duplications, an indication that the greater number of upregulated chimeras is not the product of gene sequence length (SI Appendix, Table S8). Thus, we suggest that chimeric constructs and other complex mutations that shuffle regulatory elements commonly alter expression producing immediate and drastic changes in RNA levels. In contrast, whole gene duplications rarely produce expression effects in adult gonads and soma studied here. Tandem duplications that form chimeric genes are more likely to be found at low frequency in comparison to whole gene duplications (Wilcoxon rank sum test *W* = 2452.5, *P* = 0.03881), suggesting predominantly detrimental impacts. However, chimeras have been shown to be more likely to show signals of selection favoring their spread in natural populations (11). The observed role of chimeric genes as mutations that can produce non-neutral impacts, especially in comparison to whole gene duplications, is at least partially explained by their ability to produce large magnitude changes in gene expression.

### Recruitment of non-coding sequence and *de novo* gene origination

In addition to chimeric gene structures, duplicated gene fragments that capture the 5ʹ portion of a transcript have the potential to activate neighboring sequences that were previously untranscribed, thereby creating the potential for *de novo* genes (SI Appendix, Figure S6B). We observe signs consistent with putative *de novo* gene origination through the combination of 5ʹ gene sequences with untranscribed regions during tandem duplication. We observe 43 cases of putative recruited non-coding sequence, 15 of which do not inherit a start codon from the parental gene. Among tandem duplications, we observe 30 cases associated with activation of transcription in neighboring regions that were previously untranscribed. These new genes are typically associated with duplication within a transcript or through the union of a 5ʹ UTR and neighboring non-transcribed sequence (Figure 4, Table 1). Parental genes for these cases of *de novo* gene formation include XX.

**Figure 4:**
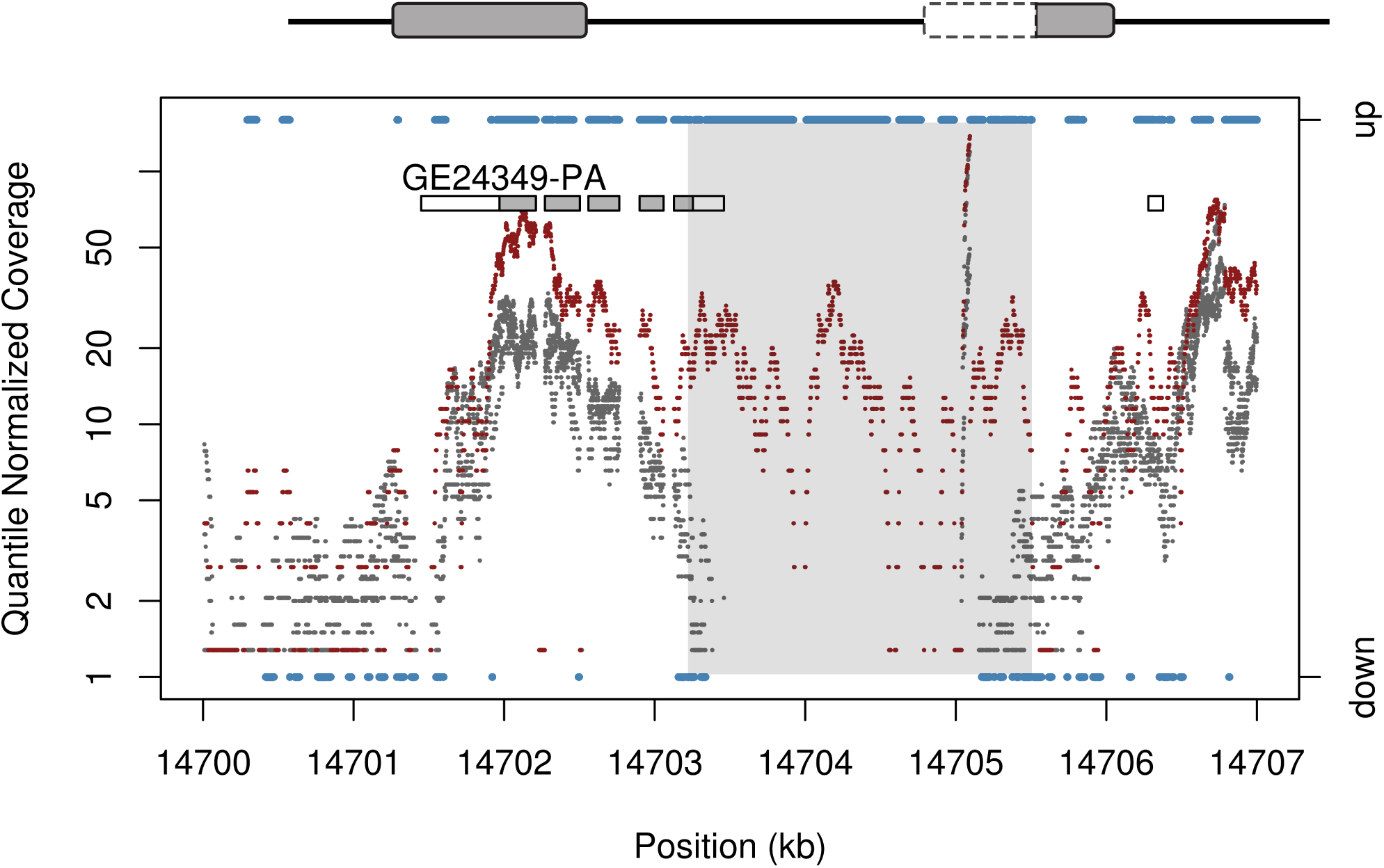
Tandem duplication creates a *de novo* gene on chromosome 3R. The 5ʹ end of GE24349 is duplicated and placed adjacent to formerly untranscribed sequence, producing transcription and putative *de novo* gene creation. The reference strain does not show transcription in the region (grey) and no other sample strain exhibits upregulated sequence across the region. Sites with upregulated or downregulated sequence as defined by HMM output is shown in blue, using the right axis. HMM state calls for sites with unchanged expression are not shown. The region spanned by the tandem duplication is shaded in grey. The tandem duplication activates a previously untranscribed region from roughly 14703500 - 14705000 bp. There is also upregulation in some exons for *GE24349*, possibly indicating a longer fusion transcript that reads through to the end of the nearest adjacent 3ʹ UTR.

In the absence of information about genome structure these will appear to be *de novo* gene creation, but with clearly defined boundaries of tandem duplications we can clarify that shuffling of 5ʹ segments of transcripts is one potential mechanism for activation of previously untranscribed regions. Among these putative cases of *de novo* activation, 23 are identified in the testes (Table 1), consistent with the out-of-the-testes hypothesis observed for new genes (29, 15). The mean size of these *de novo* expressed regions is 385 bp, with no evidence of significant size differences across tissues (*F* = 0.798, *df* = 2 *P* = 0.458; Table S9). For single transcripts, however, there can be variation in length across tissues, possibly reflecting isoform switching across tissues or general imprecision (Table S9). Reference genome expression level for parental genes that contribute to *de novo* gene formation are given in Table S10. These results offer one potential molecular mechanism to explain previously observed *de novo* gene origination, which is expected to have widespread results on evolution of new genes (30) and potential contribution to disease. Given the large number of sequences identified in such a small fraction of the genome that is spanned by tandem duplications, we would suggest that tandem duplicates can be a powerful force for new gene creation and neofunctionalization as well as contributors to pathogenic misexpression. While the predominant fate of new proto-genes is eventual loss (10, 3, 5, 31), such variants are expected to contribute a steady stream of new transcripts.

### Duplication of ancestrally carcass-expressed genes

To determine whether ancestral expression patterns of genes influence their propensity for tandem duplication, we compare genes that are captured by duplications with those that are not. Three replicates of the *D. yakuba* reference were previously assayed for differential expression across tissues (22). These reference strains contain none of the tandem duplications described here and should reflect the unmutated ancestral state. Among genes captured by duplications, 195 are biased toward ovary in the ancestral state whereas 345 are biased toward female carcass based on comparisons of ovary vs. carcass. In male somatic and germline comparisons, 168 genes captured by tandem duplication are biased toward testes in the ancestral state, and 131 are biased toward the male carcass. Based on resampling of genes in the reference, there is an excess of genes with biased expression toward female carcass (one-sided *P* < 10^−4^) and a deficit of genes that are duplicated with biased expression toward the ovaries in the ancestral state (one-sided *P* = 0.002). In males we observe an excess of genes that are duplicated with biased expression toward the carcass (one-sided *P* = 0.0029) but no bias with respect to testes expressed genes (one-sided *P* = 0.1443). Genes that duplicate have higher expression level in reference strains in every tissue (Figure 5,Table S11), pointing to the potential for biases in tandem duplicate formation or putatively selection to retain genes. Tandem duplications that are present only in 1 or 2 sample strains are expected to be newly formed, with little room for selection to bias relationships. When we limit analyses to rare variants present only in 1 or 2 sample strains, the excess of expressed genes is equally true (Table S12), suggesting that biases in formation toward transcribed regions certainly contribute to a large portion of the expression difference for duplicated sequence.

**Figure 5:**
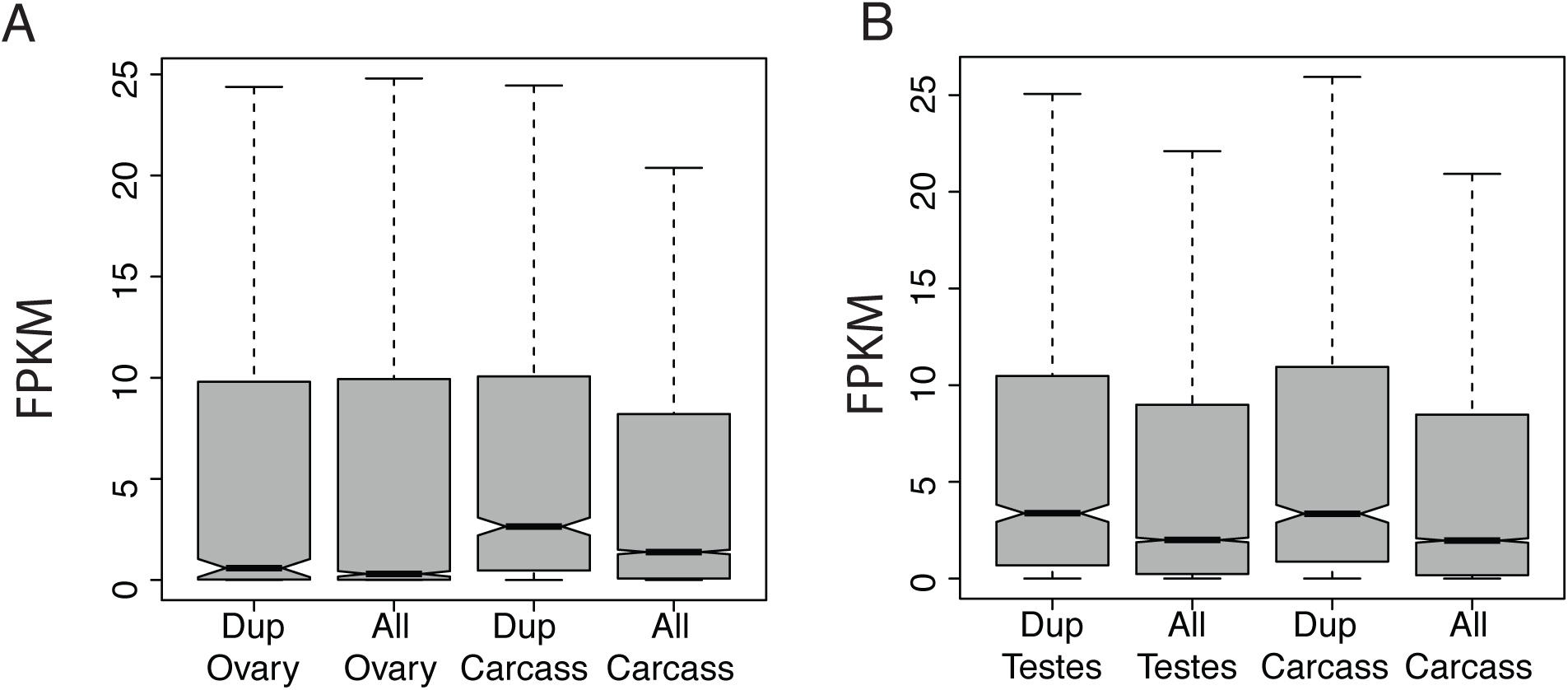
Expression levels (in FPKM) for unduplicated ancestral state for three *D. yakuba* reference replicates for genes that are duplicated in sample strains compared to expression levels for all genes. FPKM values are indicative of ancestral expression patterns prior to duplication. Duplicated genes have higher mean and median ancestral expression compared to non-duplicated genes in female tissues (A) and male tissues (B). Genes that are duplicated have lower median expression in ovary compared to carcass in females (A) but there is no difference in expression in reproductive vs. somatic tissue in males (B). Plots shown exclude outliers.

## Discussion

### Little evidence of expression differences due to whole gene duplication

One hypothesis to explain the phenotypic impacts of duplicate genes is that changes in transcript levels due to gene copy number result in novel phenotypes (7). In contrast to these common assumptions about the molecular impacts of tandem duplications, we observe little evidence for increased expression in response to duplication, with 7.6% or fewer duplicated genes showing evidence for increased expression in each tissue. These numbers are not significantly different from the random expectation based on the frequency of upregulation across the genome as a whole (Table S1). Results based on the HMM which uses site specific criteria show qualitatively similar results, with no enrichment for expression differences compared with background rates. The concordance with genome wide background rates points to the possibility of secondary mutations modifying expression or environmental effects on gene expression in spite of controlled growth conditions. Similar expression buffering has been observed in large chromosomal abnormalities in surveys for a small number of *Drosophila* mutants (32) and *Ubx* deletions often exhibit buffered phenotypes (33). The results described here suggest that these early results for small numbers of lab mutants are likely to reflect a more general genome-wide phenomenon.

The observed lack of expression changes is consistent with previous results showing that expression changes at CNVs are not commonly targets of natural selection (34). Furthermore, many such expression changes appear to be qualitative changes that are not compatible with the notion that duplication commonly results in two-fold increases in expression. The majority of genes show no evidence for asymmetrical expression of duplicates, suggesting that dosage sharing is common. These results are compatible with the hypothesis that many genes are subject to tight regulatory control and that transcription is not the limiting factor in protein production for many genes. Alternatively, it may be that promotors and full transcripts including UTRs are not sufficient to drive gene expression, implying strong cis-regulatory effects beyond the promoter. Together, these results suggest that the phenotypic impacts of tandem duplications are more complex than additive changes in transcript abundance due to copy number. Previous work has suggested that selection to maintain total expression levels across ohnologs might lead to expression subfunctionalization (35). Rather than genes increasing expression due to additive changes, then having to evolve back toward lower levels, we would suggest that genes initially are held at that same constant level through regulatory feedback loops.

Similarly low rates of expression changes for CNVs in humans (36) and rodents (37) imply that these results are likely to be general across many organisms. In humans, copy number changes are associated with a large number of diseases. For some genes, especially those where relative dosage is more likely to matter, the phenotypic and selective impacts may be different and we might expect to see different patterns for this small minority of genes (38, 24, 7). Pesticide resistance genes have been reported to have changes in gene dosage after duplication (reviewed in 7). The most highly expressed genes, which may be more likely to be transcription limited may be more likely to exhibit such expression changes from gene dosage. Indeed, recent transgenic experiments using the highly expressed gene *Adh* show transcription levels respond in response to higher copy number (39). Hemizygous deletions in *D. melanogaster* suggest that expression effects for many genes are mediated by robust regulatory architecture, but with larger effects from copy number reduction in the most highly expressed genes (40). Ohnologs, retained in the genome after whole genome duplication, also appear to be more sensitive to copy number changes than general CNVs, suggesting qualitative differences in their response to copy number (41). The whole gene duplications with upregulated expression here encompass diverse functional roles, including a testes-expressed endopeptidase, metabolism peptides, and a chorion protein. Yet, given the rarity of regulatory changes due to increases in gene copy number presented here, we suggest that alternative mechanisms are necessary to explain the role tandem duplications play in generating pathogenic phenotypes (16).

### Regulatory novelty from exon shuffling

In contrast with unaltered expression patterns among whole gene duplications, chimeric genes, UTR shuffling, and recruitment of non-coding sequence often produce changes in expression with extreme up-regulation. These variants are polymorphic, and expression effects are seen even among genes at low frequency in the sample, suggesting that many of these constructs are very young with little time to accumulate secondary mutations that might explain patterns observed. Furthermore, such changes in gene expression reflect the chimeric and fragmented gene structures produced, indicating that they are the direct product of chimera formation, not environmental effects or other spurious signals. Regulatory modules for genes can be complex, with promoters and enhancers located at 5ʹ or 3ʹ ends of genes. Additionally, transcripts may carry motifs or secondary structures that are part of regulatory feedback loops via degradation pathways (42, 43). Because chimeric genes shuffle the 5ʹ and 3ʹ ends of gene sequences, they can recombine diverse regulatory elements to generate novel expression patterns. Similarly, gain or loss of regulatory elements for gene fragments or genes that recruit non-coding sequences could produce novel combinations, resulting in altered transcript levels. Here, we observe a regulatory novelty in chimeric constructs, analogous to novel combinations of functional domains that result from exon shuffling (44, 45, 28). This regulatory novelty may explain one mechanism to generate immediate regulatory divergence between tandem duplications that can contribute to genome evolution and population level variation.

One hypothesis to explain the evolution of network structure after whole gene duplication involves loss of expression or interaction after polyploidy (46). However, we have found that upregulation, not silencing, is a common result of tandem duplication, indicating that such results reflect either major differences between polyploidy and gene expression or that present interaction and expression information does not perfectly reflect ancestral states. Previous results have suggested that duplications produce dosage changes in transcript levels(23, 12, 7). However, such results are likely the product of limited ability to detect tissue-specific changes in whole adult flies, with no tissue level resolution (for associated data description 47, 48). Separation of tissues is critical to establishing effects on gene expression, as upregulation in a single tissue that is only a fraction of the biomass will give a false signal of minor expression changes. Given the limited effect of gene copy number for whole gene duplication and the extreme expression changes associated with alternative gene structures, we suggest that such additive models of duplicate gene evolution do not reflect the full complexity of regulatory pathways or the fundamental nature of mutation.

We have observed regulatory changes and misexpression of gene fragments as a product of chimera formation, recruitment of non-coding sequence, and deletions that proceed rapidly after duplication to create variants with unusual gene structures. *De novo* proto-genes are commonly found in subtelomeric regions in yeast (31) and changes in genome structure are common in these regions as well (14) possibly explaining a portion of the pattern. One mechanism for origination of *de novo* genes that has been proposed is antisense transcription from divergent promotors (49, 50). These results offer a second mechanism that relies on canonical promoters, transcription start signals, and translation start signals with genome shuffling to serve as drivers of new gene sequences. These newly originated exons outside annotated gene sequences have a mean length of 385 bp. These are slightly shorter than previous assays of *de novo* genes (30), although these numbers do not include length of copied gene fragments.

We observe no clear evidence of divergent promoters generating new genes at the tandem duplicates surveyed here, suggesting that the two mechanisms operate independently to serve as sources of new gene sequences. Many of the *de novo* transcript sequences that are newly formed may have abnormal translation products, and most new genes that form are expected to be eventually lost (31). However, a portion of such new proto-genes can be modified by selection to form fully functional genes (31). Thus, the tandem duplications described here are expected to serve as a steady source of new gene sequences, and a minority of these are expected to be sources of novel functions (51, 31, 10, 11, 52, 53, 54, 30). RNA-seq based annotations in *D. yakuba* have identified 1340 lineage specific genes based on the *D. yakuba* reference, which do not have orthologs in other *Drosophila* genomes (22). The observed high rates of *de novo* gene formation are likely to explain a significant portion of this signal.

Previous work has found qualitatively similar results for small numbers of genes and such mutations have potential to cause other types of qualitative changes in gene regulation beyond the limited amount captured in the current study. Chimeric genes can produce differences in presence or absence of transcripts in tissues or timepoints (11, 10), and a synthetic lab-generated chimera produces differential regulation in spatial patterning of *hox* gene expression during development (9). Although differing methods of regulatory feedback mechanisms in mammals might be thought to render different effects, there are three case studies of chimeric gene formation in humans associated with expression changes, suggesting that the phenomenon deserves more careful study in human datasets. First, a chimeric gene that forces novel expression in the brain is associated with schizophrenia in humans (8). Second, a newly formed chimeric gene is known to have novel expression in human testes (55), suggesting that these results are likely to be generally applicable to studies of human health. Finally, one known case of *de novo* gene origination through chromosomal rearrangement is know to have formed a new testis-expressed gene in humans (56). Our data strongly suggest shuffling of modular genomic units can be a powerful force to develop novel regulatory profiles or unique expression patterns that has not been fully explored. We therefore suggest that these genes with altered transcription patterns are a prime source for genetic novelty, immediate neofunctionalization, and genes with widespread potential for non-neutral effects well deserving of future study in model and clinical systems.

### Mutations of major effect

Young whole gene duplications are expected to be highly similar and modification of amino acid sequences through point mutations can take many generations. Barring changes in transcript dosage, these new faithfully copied whole gene duplications are unlikely to have extreme and immediate phenotypic effects. Mutations that shuffle UTRs, recruit non-coding sequence, or combine separate coding sequence can produce regulatory changes and protein sequence changes immediately upon formation and *a priori* are more likely to produce phenotypic effects. Although many such effects are likely to be pathogenic (16, 57, 17, 20, 18, 19, 21), they may often be adaptive as well (10, 11, 52, 53, 54). Indeed, chimeric genes that combine segments of two or more coding sequences are more likely to be involved in selective sweeps immediately after formation in comparison to whole gene duplications and are a richer source of genetic novelty (11). Because many of these variants capture only portions of gene sequences (14), high-throughput use of gene models in reference strains will underreport expression differences, thereby missing a large portion of variation in gene expression that could potentially explain phenotypic variation. The use of gene-model free expression testing in high coverage data, as we have presented here, offers greater power to assay gene expression changes at abnormal gene structures and could have important impacts even in organisms outside *Drosophila*. Similar approaches can readily complement standard differential expression testing software to gain additional information in studies for the genetic basis of adaptation, quantitative genetics, and studies of pathogenic phenotypes.

We have previously described large numbers of deletions that appear rapidly after duplication (14) which here are found to be associated with expression changes. CNV identification methods that do not account for secondary deletions, or that cluster all putatively duplicated loci too broadly thereby misidentifying breakpoints will lose important information with respect to gene structure. Such missing information can have a detrimental impact on the ability to correctly identify variation, associated expression effects, and regulatory changes associated with gene fragmentation. Although common CNVs at a frequency ≥10%, which are well tagged by SNPs, are unable to explain missing complex trait and disease heritability in humans (58) the majority of tandem duplicates described here appear to be at low frequency and tandem duplicates modified by secondary deletions will be rarer still (14). Especially given the difficulties of identifying variants where linked SNPs are more common than causative mutations (59), the inability to identify modified duplicates may explain some portion of failure to identify causative variants or eQTLs in GWAS and other clinical studies (16, 18). Here, the precision that is available in *Drosophila* allows greater resolution than has been previously provided in non-model systems, allowing inferences concerning the nature of mutation that are well worth exploring in future studies of phenotype and disease in more complex genomes, including humans.

### Ancestral expression patterns of duplicated genes

We observe elevated ancestral expression level in the unduplicated reference strain for genes that are captured by duplications in at least one sample strain, suggesting that genes that are originally highly expressed are more likely to be associated with duplications (Figure 5, Table S11). Even limiting the genes surveyed to genes that are identified in only one or two strains, expression still appears to be elevated above the genome wide background (Table S11). Thus, we suggest that genes that duplicate are more likely to be expressed or are more highly expressed in the unduplicated ancestral state compared to the genome wide average. This pattern is observed in male and female somatic and reproductive tissues as well as low-frequency variants, making it unlikely that selection on a single functional category or gene family is responsible for the duplication of transcribed genes.

Tandem duplications can form through several mechanisms, including replication slippage, ectopic recombination, aberrant DNA break repair, and non-homologous end joining. Transcription-coupled repair and the avoidance of repair in regions bound by nucleosomes is commonly invoked to explain mutational patterns for SNPs in mammals and yeast (60, 61). However there is no strong evidence for such transcript coupled repair in *Drosophila* (62, 63). Genes that are transcribed are often members of open chromatin, and it is possible that the correlation between actively transcribed genes and chromatin states might promote greater recombination and repair and thereby explain the excess of transcribed genes among tandem duplications. We observe equal levels of upregulation for chimeric gene segments in female germline as in male germline, but lower fold-change in the testes (Figure 1). Because many genes are already expressed in the testes, chimeric portions which are already highly expressed are less likely to show high level upregulation under a scheme of non-additive expression effects from shuffling of regulatory elements. Similarly, widespread transcription of parental genes in the ancestral state rather than selection is likely to explain the overabundance of novel gene expression we observe in the testes due to a simple abundance of testes-driving promoters. This widespread transcription may be due to spurious, non-functional transcription in the testes, which combined with tandem duplication can be a fortuitous but powerful source of new genes.

## Methods

### Identifying tandem duplications and gene expression changes

We identified tandem duplications using paired-end Illumina genomic sequencing, as previously described (14). Briefly, tandem duplications were defined by three or more divergently oriented read pairs that lie within 25 kb of one another. We excluded duplications indicated with divergent read pairs in the reference strain, which are indicative of technical challenges or reference mis-assembly. We also excluded duplicates which were present in *D. erecta*, resulting in a high quality data set of newly derived tandem duplications that are segregating in natural populations. Duplications were clustered across strains within a threshold distance of 200 bp and the maximum span of divergently oriented reads across all strains were used to define the span of each duplication. We then identified gene sequences captured by tandem duplications using RNA-seq based gene models previously described in Rogers et al (22).

RNA-seq samples were prepared from virgin flies collected within 2 hrs. of eclosion, then aged 2-5 days post eclosion before dissection. We dissected ovaries and headless carcass for adult females, and testes plus glands for adult males. Samples were flash frozen in liquid nitrogen and stored at −80°C before extraction in trizol. Illumina sequencing libraries were prepared using the Nextrera library preparation kit, and were sequenced on an Illumina HiSeq 2500. Fastq data were aligned to the *D. yakuba* reference genome using Tophat v.2.0.6 and Bowtie2 v.2.0.2 (64). Site specific changes in gene expression were determined using a Hidden Markov Model that implements the underlying statistical model of the Cufflinks suite (25). Further description of RNA-seq sample preparation, data analysis, and HMM performance is available in SI Appendix. Sequence data are available in the NCBI SRA under PRJNA269314 and PRJNA196536. Code is available at https://github.com/evolscientist/ExpressionHMM.git.

### Sample preparation and RNA-sequencing

We gathered RNA-seq data for 15 samples and the reference genome (Table S13). Fly stocks were incubated under controlled conditions at 25°C and 40% humidity. Virgin flies were collected within 2 hrs. of eclosion, then aged 2-5 days post eclosion before dissection. We dissected samples in isotonic Ringers solution, using female ovaries and headless gonadectomized carcass from two adult flies as well as testes plus glands and male headless gonadectomized carcass for four adult flies for each sample RNA prep. We collected three biological replicates of the *D. yakuba* reference, and one replicate per sample strain for 15 samples of *D. yakuba*. Samples were flash frozen in liquid nitrogen immediately after dissection, and and stored in 0.2ml Trizol at −80°C. All samples were homogenized in 0.5ml Trizol Reagent (Invitrogen) with plastic pestle in 1.5ml tube, mixed with 0.1ml chloroform, and centrifuged 12,000g 15min at 4oC, as Trizol RNA extraction protocol. The RNAs in the supernatant about 0.4ml were then collected and purified with Direct-Zol RNA MiniPrep Kit (Zymo), followed the protocol. The total RNAs were eluted in 65*μ*L RNase-Free H_2_O. About 1*μ*g purified RNAs were treated with 2*μ*L Turbo DNase (Invitrogen) in 65*μ*L reaction, incubated 15min at room temperature with gentle shaking. These RNAs were further purified with RNA Clean and Concentrator-5 (Zymo). One extra wash with fresh 80% ethanol after the final wash step was added into the original protocol. The treated RNAs were eluted with 15*μ*L RNAse-Free H_2_O, and stored at −80°C.

The amplified cDNAs were prepared from 100ng DNase treated RNA with Ovation RNA-Seq System V2 (Nugen) and modified protocol. The preparations followed the protocol to the step of SPIA Amplification (Single Primer Isothermal Amplification). The amplified cDNAs were first purified with Purelink PCR Purification Kit (Invitrogen, HC Binding Buffer) and eluted in 100*μ*L EB (Invitrogen). These cDNAs were purified again to 25*μ*L EB with DNA Clean and Concentrator −5 Kit (Zymo) for Nextera library preparation. About 43ng cDNAs were used to construct libraries with Nextera DNA Sample Preparation Kit (Illumina) and modified protocol. After Tagmentation, Purelink PCR Purification Kit with HC Binding Buffer was used for purification and eluted with 30*μ*L EB or H_2_O. The products (libraries) of final PCR amplification were purified with DNA Clean and Concentractor-5 and eluted in 20*μ*L EB. The average library lengths roughly 500bp were estimated from profiles of Bioanalyzer (Agilent) with DNA HS Assay. All libraries were normalized to 2-10nM based on real-time PCR method with Kapa Library Quant Kits (Kapa Biosystems). The qualities and quantities of these RNAs, cDNAs and final libraries were measured from Bioanalyzer with RNA HS or DNA HS Assays and Qubit (Invitrogen) with RNA HS or DNA HS Reagents, respectively. Samples were barcoded and sequenced in 4-plex with 76 bp reads on an Illumina HiSeq 2500 using standard Illumina barcodes, resulting in high coverage with thousands of reads for *Adh*, the most highly expressed gene in *Drosophila* (Figure S7). We sequenced one replicate per sample strain as well as three biological replicates of each reference strain for all tissues. Female tissues for sample strains and one replicate of the reference genome were sequenced with single end reads, while two replicates of reference genome female tissues and all male tissue samples were sequenced with paired end reads.

### Reference expression patterns

Expression patterns in the reference genome, indicative of the ancestral, unduplicated state, were established according to Rogers et al. (22). Briefly, sequences were mapped to the genome using Tophat v.2.0.6 and Bowtie2 v.2.0.2, using reference annotations as a guide, ignoring reads which fell outside reference annotations (-G). We estimated transcript abundances and tested for differential expression at an FDR ≤ 0.1 using Cuffdiff from Cufflinks v. 2.0.2 with quantile normalized expression values (-N), again using only reads which aligned to annotated gene sequences. All other parameters were set to default. We compared female ovaries to female carcass and male testes to male carcass for the reference strain replicates to determine tissue biased expression prior to duplication. Overrepresentation and underrepresentation of genes with tissue biased expression were established by resampling 10,000 replicates of randomly selected genes.

### Duplicated gene sequences

We used gene models developed from RNA-seq guided reannotation of the *D. yakuba* reference genome (22). The maximum span of divergently oriented reads was considered the bounds of duplication, similar to previous analysis (14) using FlyBase gene models (13). These revised gene models include 5ʹ and 3ʹ UTRs, and are essential to correctly establish the effects tandem duplicates will have on gene structures. These revised gene models show greater concordance with *D. melanogaster*, resulting in an additional 1000 *D. melanogaster* genes with an ortholog in *D. yakuba* compared to previous gene annotations (22). We additionally identify 1340 lineage specific genes in *D. yakuba*, hundreds of which display expression bias across tissues (22).

### Differential expression testing using cuffdiff

Sequences for each reference replicate and barcoded sample strain were mapped to the genome using Tophat v.2.0.6 and Bowtie2 v.2.0.2, using reference annotations (22) as a guide on the *D. yakuba* r1.3 reference genome, ignoring reads which fell outside reference annotations (-G). We estimated transcript abundances and tested for differential expression in an all-by-all comparison at an FDR ≤ 0.1 using Cuffdiff from Cufflinks v. 2.0.2 with quantile normalized expression values (-N), again using only reads which aligned to annotated gene sequences with all other parameters set to default. Reference replicates were grouped for differential expression testing in Cuffdiff. For each tissue the total number of duplications displaying increases in expression for whole gene duplication and for background rates were compared using a chi-squared test with 1 degree of freedom.

### Test of dosage-sharing

One hypothesis for the lack of gene expression changes among whole gene duplications is that secondary mutations might result in asymmetric silencing of one duplicate copy. If duplicate copies have differentiated from one another, this should be apparent in large numbers of seemingly heterozygous sites in the genomic SNP data. To test for differential expression among copies of whole gene duplication, we identified all putatively ‘heterozygous’ sites that might indicate differentiating SNPs across copies. Using samtools mpileup (v. 1.3) and bcftools consensus caller (v.1.3) with parameters set to default, we identified all putatively heterozygous sites in the genomic sequences for each strain. We then generated SNP calls using identical criteria for RNA sequencing data. The number of reads supporting heterozygous calls for the reference sequence and SNP sequence were then compared using a Fisher’s exact test. Only SNPs with at least 10 reads covering the site in both genomic and RNA sequencing datasets were used for differential expression testing. Sites which exhibited significant differential expression of SNPs in at least one strain that housed a duplication were considered candidates for differential expression of duplicate copies. Similar signals could be produced by allele specific expression even at unduplicated sites. We filtered out all sites that displayed such allele specific expression in strains that did not contain the duplication in question, as these are unlikely to reflect processes specific the duplication.

### HMM for expression patterns

Coverage in mapped RNA-seq data per site for each strain was calculated using samtools depth. Sample strains show variable FPKM based on cuffdiff analysis (Figure S8-S9), which might potentially influence power to detect differential expression. To reduce the influence of coverage differences across samples and generate more robust expression calls (65), we quantile normalized each chromosome in R so that coverage per site across all strains has the same mean and variance for a given chromosome in a given tissue. Mean quantile-normalized coverage among regions corresponding to annotated exon sequences was 61 X. This quantile normalized coverage depth per site was used as input for a Hidden Markov Model (HMM) to identify site specific changes in gene expression, offering differential expression testing independent of gene models and exon annotations. This gene-model free expression testing is essential for discovering the regulatory impacts of complex mutations such as chimeric genes, recruited non-coding sequence, and duplication-deletion constructs all of which do not respect gene boundaries. This HMM also performs comparative hypothesis testing, choosing the most likely expression state for each site, rather than simply testing adherence to a null statistical model, an important methodological advantage.

The HMM attempts to identify three underlying states: decreased expression, stable expression, and increased expression. Initial state probabilities were set according to *π*_0_ and transition probabilities were set according to *T*, where row and column indices 0,1,2 are indicative of decreased, stable, and increased expression, respectively. Initial probabilities are set such that the singleton state is initially most likely and states are initially most likely to remain constant during transitions.

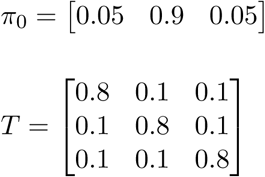

Very low transition probabilities can have a chilling effect on output of HMMs, which might potentially bias results away from detecting expression changes, a major hypothesis that is tested in the current work. However, results with alternate transition matrices defined by the Baum-Welch algorithm do not differ qualitatively from those presented in the main text (Table S14). This is equally true for *de novo* genes.

Emission probabilities were modeled as follows: We compare the ratio of quantile normalized coverage per site for each sample strain to the mean for the three reference replicates. We assume the natural log of the fold change is normally distributed. Under a null model of no expression change, we can assume mean and variance in the sample will be equal to the mean and variance in the reference replicates, and use the delta method to approximate the variance, a common method of variance estimation in differential expression testing (25). Under such an approximation, the variance of the natural log of the fold change is equal to 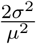 where *σ*^2^ is the observed variance in quantile normalized coverage for the reference variance and *μ* is the observed mean quantile normalized coverage in the reference replicates. For stable expression, the distribution of the natural log of the mean fold change should be centered about 1, corresponding to no expression difference.

For increased expression we again assume a normal distribution for the log fold change, but assuming a true mean quantile normalized coverage at the upper critical value of the distribution under no difference in gene expression. For decreased expression we again model the log fold change as a normal distribution, but assume a true mean of quantile normalized coverage at the lower critical value of the distribution under no difference in gene expression. We model the likelihood of the data given no change in expression as the probability of a test statistic with an absolute value as large or larger than the observed, given a normal distribution of the log mean fold change. For sites with increased expression, we model emission probabilities as the probability of a test statistic at least as high as that observed. For sites with decreased expression, we model emission probabilities the probability of a test statistic at least as low as that observed.

The log fold-change distribution for emission probabilities is unable to accurately assign likelihood of upregulated expression if the mean coverage in all reference strains is close to zero. In cases where the reference strain mean for three replicates was less than 0.5, if sample strains exhibited coverage greater than 5 or more reads, we assigned a probability of upregulation of 0.95 as these indicate clear signs of upregulation of silenced sequence, but otherwise assigned a probability of stable expression of 0.95. State decoding was performed using the Forward-Backward algorithm, which maximizes the number of correctly predicted states (66). The choice to maximize predictions per site rather than the most likely path (using the Viterbi algorithm) is important to maintain decoding of independent results across sites given the use of the HMM in site-specific differential expression testing. The use of high coverage RNA-seq data is essential for accurate performance of the HMM to detect site specific changes in expression and applications in lower coverage sequencing may have reduced power. Plots of HMM output with quantile normalized RNA-seq data show that the HMM detects increased and decreased expression for modest expression differences (Figure S4).

For each chimeric gene and whole gene duplication, we used the HMM output by tissue to define genes where duplicated sequence has been significantly upregulated in response to tandem duplication. We require that each gene or gene fragment have at least 50% of annotated exon sequence upregulated, considering only blocks of upregulated sequence 50 bp or longer. For putative cases of *de novo* gene creation, we identified blocks of upregulated sequence 50 bp or longer which do not overlap with annotated exons, and which do not have quantile normalized coverage above 2.0 in the three reference replicates. We then retained only cases that spanned at least 200 bp of the tandem duplication, in accordance with methods used by Zhao et al. (30). Performance of the HMM to call sites with increased and decreased expression is shown in Figure S4. Genes with signals of expression changes in at least one strain were considered to be upregulated.

### Mean fold change comparisons

To further establish regulatory profiles for each chimeric gene and whole gene duplication, we additionally estimated the mean fold change across all sites. This data are independent of HMM performance and gives a detailed portrait of the quantile normalized coverage data. We estimate mean coverage per site across all sites in sample and reference for a given chimera segment in a given strain. We consider segments independently as parental genes may have differing levels of ancestral expression in the reference strain. The ratio of mean coverage in the sample to mean coverage in the reference is then recorded as mean fold change per site, placing a lower bound on reference coverage level of one read per site. The mean fold change for each chimeric gene and each duplicate gene is plotted in Figure 1. The mean fold change for chimeric genes were compared to the mean fold change at the same gene fragments in strains that lacked the duplication in question in individual tissues using a Wilcoxon rank sum test.

## Acknowledgements

We would like to thank Rahul Warrior for providing temperature and humidity controlled incubator space and Rachel Martin for an emergency source of liquid nitrogen. Andrew Foran provided technical assistance.

RLR, LS, and KRT designed experiments; RLR and LS collected data; RLR developed new analytical tools; RLR and KRT performed analyses and wrote the paper. We also thank our anonymous reviewers whose comments substantially improved the manuscript.

## Supporting Information

DuplicationCoordsReFmt.txt - Duplications

DupTransCoordsReFmt.txt - Duplicated gene sequences

MutationTypes.txt - Chimeric Genes, Recruited non-coding sequences, and Whole Gene Duplications

README.txt - Readme file

RecruitNonCoding.GO.txt - Information on recruited non-coding sequences

ReadsPerGene.carcass.txt - Reads per gene for female carcass

ReadsPerGene.ova.txt - Reads per gene for female ovary

ReadsPerGene.malecar.txt - Reads per gene for male carcass

ReadsPerGene.testes.txt - Reads per gene for male testes

FigS1.pdf - Figure S1

FigS2.pdf - Figure S2

FigS3.pdf - Figure S3

FigS4.pdf - Figure S4

FigS5.pdf - Figure S5

FigS6.pdf - Figure S6

FigS7.pdf - Figure S7

FigS8.pdf - Figure S8

FigS9.pdf - Figure S9

TableS1.pdf - Table S1

TableS2.pdf - Table S2

TableS3.pdf - Table S3

TableS4.pdf - Table S4

TableS5.pdf - Table S5

TableS6.pdf - Table S6

TableS7.pdf - Table S7

TableS8.pdf - Table S8

TableS9.pdf - Table S9

TableS10.pdf - Table S10

TableS11.pdf - Table S11

TableS12.pdf - Table S12

TableS13.pdf - Table S13

TableS14.pdf - Table S14

---

**Table S1:** Genes upregulated using cuffdiff by tissue

| Tissue | Duplicates Upregulated | Assayed | Background Upregulated | Assayed | $\chi^2$ | *P*-value |
| --- | --- | --- | --- | --- | --- | --- |
| Male Carcass | 4 | 52 | 1861 | 13174 | 0.9697 | 0.3268 |
| Male Testes | 3 | 52 | 1375 | 13174 | 0.6097 | 0.4349 |
| Female Carcass | 4 | 52 | 1733 | 13174 | 0.6993 | 0.4030 |
| Female Ovary | 4 | 52 | 1343 | 13174 | 0.0977 | 0.7546 |

**Table S2:** Whole gene duplications with upregulated expression using Cuffdiff

| tissue | gene | strain | Ref FPKM | Sample FPKM | corrected $P$-value |
| --- | --- | --- | --- | --- | --- |
| Male Carcass | 2.g417 | line9 | 74.8960 | 275.4010 | 0.00102288 |
| | GE11098 | line9 | 36.3276 | 200.8140 | $4.0626 \times 10^{-6}$ |
|  | GE11098 | line10 | 36.3276 | 132.5500 | 0.00133 |
|  | GE24648 | line1 | 2.7300 | 9.2186 | 0.0260 |
|  | GE26061 | line9 | 10.9718 | 28.8023 | 0.0371 |
| Male Testes | 2.g556 | line5 | 0.3080 | 2.27934 | 0.0122 |
|  | 2.g556 | line15 | 0.3080 | 2.0057 | 0.0230 |
| | GE11098 | line10 | 620.5090 | 4025.7000 | $9.7604 \times 10^{-4}$ |
|  | GE14157 | line13 | 1.1743 | 8.2437 | 0.0030 |
| Female Carcass | GE13159 | line19 | 2.0217 | 7.96624 | 0.0031 |
|  | GE14157 | line2 | 0.6146 | 2.8779 | 0.0046 |
| | GE20775 | line8 | 1.6645 | 18.4707 | $2.20709 \times 10^{-5}$ |
|  | GE26133 | line14 | 22.0725 | 58.6217 | 0.0421 |
| Female Ovary | 2.g556 | line2 | 0.0686 | 0.7511 | 0.0019 |
| | 2.g556 | line5 | 0.0686 | 2.6958 | $1.13228 \times 10^{-8}$ |
| | 2.g556 | line6 | 0.0686 | 1.1139 | $9.73427 \times 10^{-5}$ |
| | 2.g556 | line9 | 0.0686 | 2.0189 | $3.16552 \times 10^{-7}$ |
|  | 2.g556 | line13 | 0.0686 | 8.0706 | 0 |
| | 2.g556 | line15 | 0.0686 | 4.6893 | $6.89843 \times 10^{-12}$ |
| | 2.g556 | line19 | 0.0686 | 1.8320 | $8.96617 \times 10^{-7}$ |
| | GE24648 | line1 | 5.2891 | 28.7428 | $1.96519 \times 10^{-7}$ |
|  | GE26061 | line9 | 8.1007 | 24.2368 | 0.0040 |
| | GE24030 | line6 | 2.0279 | 10.5760 | $1.17696 \times 10^{-5}$ |

**Table S3:**
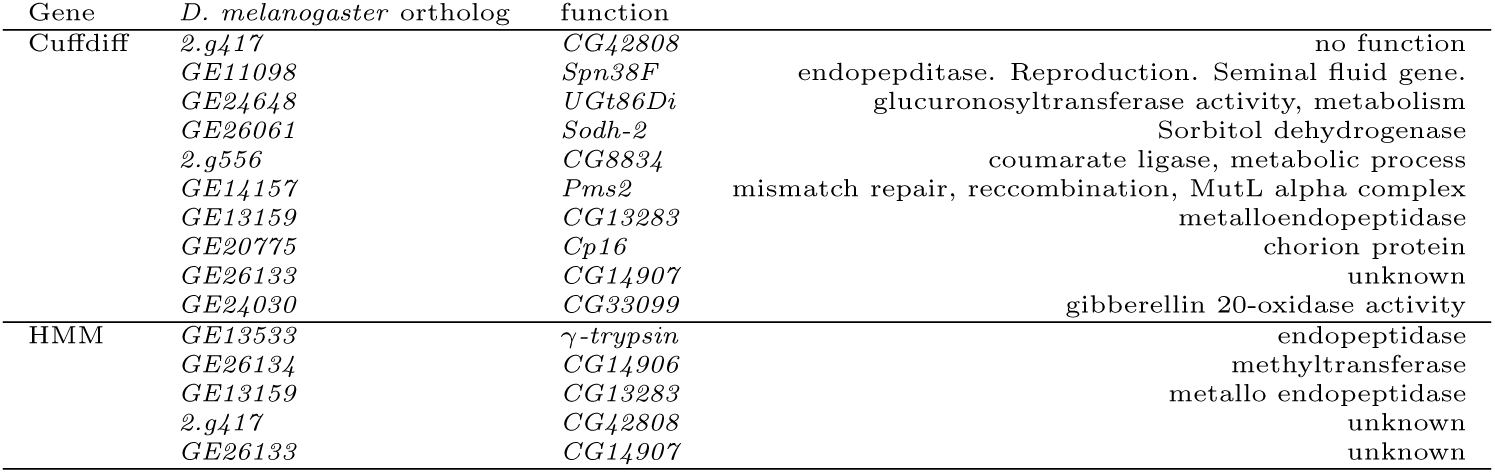
Functions of whole gene duplications with upregulated expression

**Table S4:** Genes upregulated using cuffdiff tissue, singleton variants only

| Tissue | Duplicates Upregulated | Assayed | Background Upregulated | Assayed | $\chi^2$ (2 *df*) | *P*-value |
| --- | --- | --- | --- | --- | --- | --- |
| Male Carcass | 2 | 33 | 1861 | 13174 | 0.8821 | 0.3476 |
| Male Testes | 0 | 33 | 1375 | 13174 | 2.4248 | 0.1194 |
| Female Carcass | 2 | 33 | 1733 | 13174 | 0.6826 | 0.4087 |
| Female Ovary | 2 | 33 | 1343 | 13174 | 0.1844 | 0.6676 |

**Table S5:**
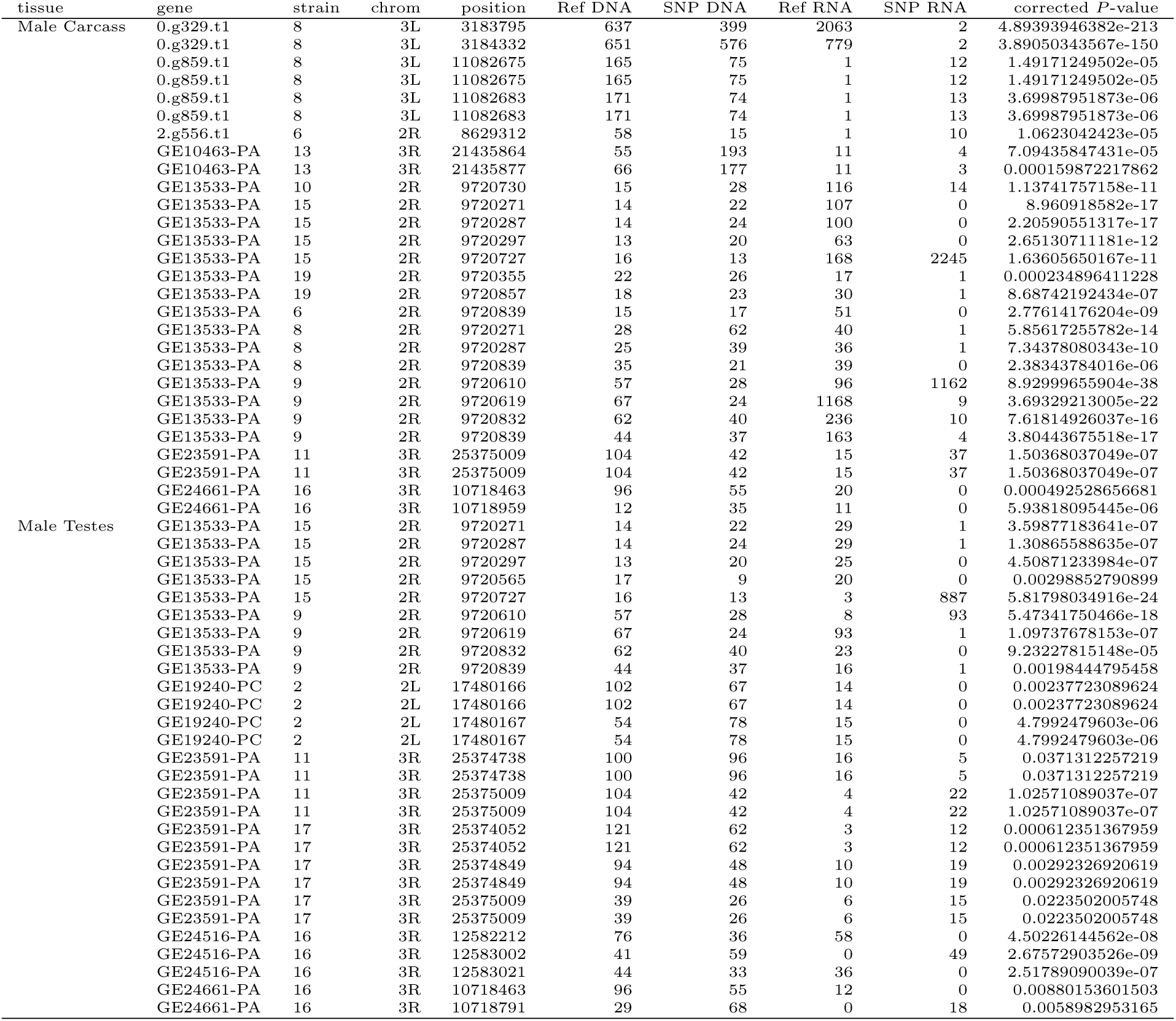
SNPs in whole gene duplications with significantly asymmetric expression in male tissues

| tissue | gene | strain | chrom | position | Ref DNA | SNP DNA | Ref RNA | SNP RNA | corrected *P*-value |
| --- | --- | --- | --- | --- | --- | --- | --- | --- | --- |
| Male Carcass | 0.g329.t1 | 8 | 3L | 3183795 | 637 | 399 | 2063 | 2 | 4.89393946382e-213 |
|  | 0.g329.t1 | 8 | 3L | 3184332 | 651 | 576 | 779 | 2 | 3.89050343567e-150 |
|  | 0.g859.t1 | 8 | 3L | 11082675 | 165 | 75 | 1 | 12 | 1.49171249502e-05 |
|  | 0.g859.t1 | 8 | 3L | 11082675 | 165 | 75 | 1 | 12 | 1.49171249502e-05 |
|  | 0.g859.t1 | 8 | 3L | 11082683 | 171 | 74 | 1 | 13 | 3.69987951873e-06 |
|  | 0.g859.t1 | 8 | 3L | 11082683 | 171 | 74 | 1 | 13 | 3.69987951873e-06 |
|  | 2.g556.t1 | 6 | 2R | 8629312 | 58 | 15 | 1 | 10 | 1.0623042423e-05 |
|  | GE10463-PA | 13 | 3R | 21435864 | 55 | 193 | 11 | 4 | 7.09435847431e-05 |
|  | GE10463-PA | 13 | 3R | 21435877 | 66 | 177 | 11 | 3 | 0.000159872217862 |
|  | GE13533-PA | 10 | 2R | 9720730 | 15 | 28 | 116 | 14 | 1.13741757158e-11 |
|  | GE13533-PA | 15 | 2R | 9720271 | 14 | 22 | 107 | 0 | 8.960918582e-17 |
|  | GE13533-PA | 15 | 2R | 9720287 | 14 | 24 | 100 | 0 | 2.20590551317e-17 |
|  | GE13533-PA | 15 | 2R | 9720297 | 13 | 20 | 63 | 0 | 2.65130711181e-12 |
|  | GE13533-PA | 15 | 2R | 9720727 | 16 | 13 | 168 | 2245 | 1.63605650167e-11 |
|  | GE13533-PA | 19 | 2R | 9720355 | 22 | 26 | 17 | 1 | 0.000234896411228 |
|  | GE13533-PA | 19 | 2R | 9720857 | 18 | 23 | 30 | 1 | 8.68742192434e-07 |
|  | GE13533-PA | 6 | 2R | 9720839 | 15 | 17 | 51 | 0 | 2.77614176204e-09 |
|  | GE13533-PA | 8 | 2R | 9720271 | 28 | 62 | 40 | 1 | 5.85617255782e-14 |
|  | GE13533-PA | 8 | 2R | 9720287 | 25 | 39 | 36 | 1 | 7.34378080343e-10 |
|  | GE13533-PA | 8 | 2R | 9720839 | 35 | 21 | 39 | 0 | 2.38343784016e-06 |
|  | GE13533-PA | 9 | 2R | 9720610 | 57 | 28 | 96 | 1162 | 8.92999655904e-38 |
|  | GE13533-PA | 9 | 2R | 9720619 | 67 | 24 | 1168 | 9 | 3.69329213005e-22 |
|  | GE13533-PA | 9 | 2R | 9720832 | 62 | 40 | 236 | 10 | 7.61814926037e-16 |
|  | GE13533-PA | 9 | 2R | 9720839 | 44 | 37 | 163 | 4 | 3.80443675518e-17 |
|  | GE23591-PA | 11 | 3R | 25375009 | 104 | 42 | 15 | 37 | 1.50368037049e-07 |
|  | GE23591-PA | 11 | 3R | 25375009 | 104 | 42 | 15 | 37 | 1.50368037049e-07 |
|  | GE24661-PA | 16 | 3R | 10718463 | 96 | 55 | 20 | 0 | 0.000492528656681 |
|  | GE24661-PA | 16 | 3R | 10718959 | 12 | 35 | 11 | 0 | 5.93818095445e-06 |
| Male Testes | GE13533-PA | 15 | 2R | 9720271 | 14 | 22 | 29 | 1 | 3.59877183641e-07 |
|  | GE13533-PA | 15 | 2R | 9720287 | 14 | 24 | 29 | 1 | 1.30865588635e-07 |
|  | GE13533-PA | 15 | 2R | 9720297 | 13 | 20 | 25 | 0 | 4.50871233984e-07 |
|  | GE13533-PA | 15 | 2R | 9720565 | 17 | 9 | 20 | 0 | 0.00298852790899 |
|  | GE13533-PA | 15 | 2R | 9720727 | 16 | 13 | 3 | 887 | 5.81798034916e-24 |
|  | GE13533-PA | 9 | 2R | 9720610 | 57 | 28 | 8 | 93 | 5.47341750466e-18 |
|  | GE13533-PA | 9 | 2R | 9720619 | 67 | 24 | 93 | 1 | 1.09737678153e-07 |
|  | GE13533-PA | 9 | 2R | 9720832 | 62 | 40 | 23 | 0 | 9.23227815148e-05 |
|  | GE13533-PA | 9 | 2R | 9720839 | 44 | 37 | 16 | 1 | 0.00198444795458 |
|  | GE19240-PC | 2 | 2L | 17480166 | 102 | 67 | 14 | 0 | 0.00237723089624 |
|  | GE19240-PC | 2 | 2L | 17480166 | 102 | 67 | 14 | 0 | 0.00237723089624 |
|  | GE19240-PC | 2 | 2L | 17480167 | 54 | 78 | 15 | 0 | 4.7992479603e-06 |
|  | GE19240-PC | 2 | 2L | 17480167 | 54 | 78 | 15 | 0 | 4.7992479603e-06 |
|  | GE23591-PA | 11 | 3R | 25374738 | 100 | 96 | 16 | 5 | 0.0371312257219 |
|  | GE23591-PA | 11 | 3R | 25374738 | 100 | 96 | 16 | 5 | 0.0371312257219 |
|  | GE23591-PA | 11 | 3R | 25375009 | 104 | 42 | 4 | 22 | 1.02571089037e-07 |
|  | GE23591-PA | 11 | 3R | 25375009 | 104 | 42 | 4 | 22 | 1.02571089037e-07 |
|  | GE23591-PA | 17 | 3R | 25374052 | 121 | 62 | 3 | 12 | 0.000612351367959 |
|  | GE23591-PA | 17 | 3R | 25374052 | 121 | 62 | 3 | 12 | 0.000612351367959 |
|  | GE23591-PA | 17 | 3R | 25374849 | 94 | 48 | 10 | 19 | 0.00292326920619 |
|  | GE23591-PA | 17 | 3R | 25374849 | 94 | 48 | 10 | 19 | 0.00292326920619 |
|  | GE23591-PA | 17 | 3R | 25375009 | 39 | 26 | 6 | 15 | 0.0223502005748 |
|  | GE23591-PA | 17 | 3R | 25375009 | 39 | 26 | 6 | 15 | 0.0223502005748 |
|  | GE24516-PA | 16 | 3R | 12582212 | 76 | 36 | 58 | 0 | 4.50226144562e-08 |
|  | GE24516-PA | 16 | 3R | 12583002 | 41 | 59 | 0 | 49 | 2.67572903526e-09 |
|  | GE24516-PA | 16 | 3R | 12583021 | 44 | 33 | 36 | 0 | 2.51789090039e-07 |
|  | GE24661-PA | 16 | 3R | 10718463 | 96 | 55 | 12 | 0 | 0.00880153601503 |
|  | GE24661-PA | 16 | 3R | 10718791 | 29 | 68 | 0 | 18 | 0.0058982953165 |

**Table S6:** SNPs in whole gene duplications with significantly asymmetric expression in female tissues

| tissue | gene | strain | chrom | position | Ref Genomic | SNP Genomic | Ref RNA-seq | SNP RNA-seq | corrected *P*-value |
| --- | --- | --- | --- | --- | --- | --- | --- | --- | --- |
| Female Carcass | 2.g556.t1 | 9 | 2R | 8628170 | 52 | 61 | 13 | 0 | 0.000134635282947 |
|  | GE13533-PA | 2 | 2R | 9720296 | 7 | 21 | 249 | 2 | 1.37883018113e-23 |
|  | GE13533-PA | 2 | 2R | 9720297 | 7 | 22 | 248 | 0 | 7.61164181427e-27 |
|  | GE13533-PA | 6 | 2R | 9720271 | 21 | 11 | 38 | 0 | 5.9627472227e-05 |
|  | GE13533-PA | 6 | 2R | 9720287 | 16 | 10 | 38 | 0 | 3.50671569654e-05 |
|  | GE13533-PA | 6 | 2R | 9720839 | 15 | 17 | 39 | 0 | 5.46168656421e-08 |
|  | GE13533-PA | 8 | 2R | 9720271 | 28 | 62 | 60 | 0 | 1.73436067901e-20 |
|  | GE13533-PA | 8 | 2R | 9720287 | 25 | 39 | 58 | 0 | 4.23964503844e-15 |
|  | GE13533-PA | 8 | 2R | 9720839 | 35 | 21 | 5647 | 6 | 2.64131038422e-39 |
|  | GE13533-PA | 9 | 2R | 9720610 | 57 | 28 | 67 | 1214 | 1.36030183298e-44 |
|  | GE13533-PA | 9 | 2R | 9720619 | 67 | 24 | 1177 | 2 | 4.3505030628e-27 |
|  | GE13533-PA | 9 | 2R | 9720832 | 62 | 40 | 619 | 12 | 1.53640332343e-27 |
|  | GE13533-PA | 9 | 2R | 9720839 | 44 | 37 | 335 | 1 | 4.32719602547e-29 |
|  | GE13533-PA | 10 | 2R | 9720730 | 15 | 28 | 468 | 4 | 3.64633793886e-31 |
|  | GE13533-PA | 15 | 2R | 9720271 | 14 | 22 | 445 | 0 | 6.83413786614e-29 |
|  | GE13533-PA | 15 | 2R | 9720287 | 14 | 24 | 438 | 1 | 1.36160863106e-29 |
|  | GE13533-PA | 15 | 2R | 9720297 | 13 | 20 | 294 | 0 | 1.29111129899e-23 |
|  | GE13533-PA | 15 | 2R | 9720727 | 16 | 13 | 658 | 1 | 8.38049203847e-19 |
|  | GE13533-PA | 19 | 2R | 9720355 | 22 | 26 | 48 | 1 | 1.91720088454e-09 |
|  | GE13533-PA | 19 | 2R | 9720857 | 18 | 23 | 41 | 2 | 1.44452774476e-07 |
|  | GE13533-PA | 19 | 2R | 9720875 | 20 | 38 | 0 | 32 | 4.15947174032e-05 |
|  | GE20775-PA | 8 | 3L | 3185046 | 570 | 372 | 25 | 0 | 7.36488931564e-06 |
|  | GE20775-PA | 8 | 3L | 3185052 | 536 | 357 | 25 | 0 | 7.26234794398e-06 |
|  | GE20775-PA | 8 | 3L | 3185112 | 377 | 517 | 22 | 0 | 8.20383183941e-09 |
|  | GE20775-PA | 8 | 3L | 3185229 | 178 | 162 | 18 | 0 | 2.27731906792e-05 |
| Female Ovary | 0.g329.t1 | 8 | 3L | 3183371 | 300 | 246 | 24 | 0 | 1.43200054203e-06 |
|  | 0.g329.t1 | 8 | 3L | 3183545 | 524 | 329 | 29 | 0 | 1.44529786542e-06 |
|  | GE19240-PC | 2 | 2L | 17480167 | 54 | 78 | 14 | 0 | 1.0224484785e-05 |
|  | GE19240-PC | 2 | 2L | 17480167 | 54 | 78 | 14 | 0 | 1.0224484785e-05 |
|  | GE21202-PA | 9 | 3L | 1897080 | 96 | 42 | 47 | 0 | 1.04727614702e-06 |
|  | GE24516-PA | 16 | 3R | 12582212 | 76 | 36 | 247 | 4 | 1.43363038229e-16 |
|  | GE24516-PA | 16 | 3R | 12583002 | 41 | 59 | 31 | 158 | 7.49321729398e-06 |
|  | GE24516-PA | 16 | 3R | 12583021 | 44 | 33 | 155 | 24 | 9.30200258675e-07 |

**Table S7:**
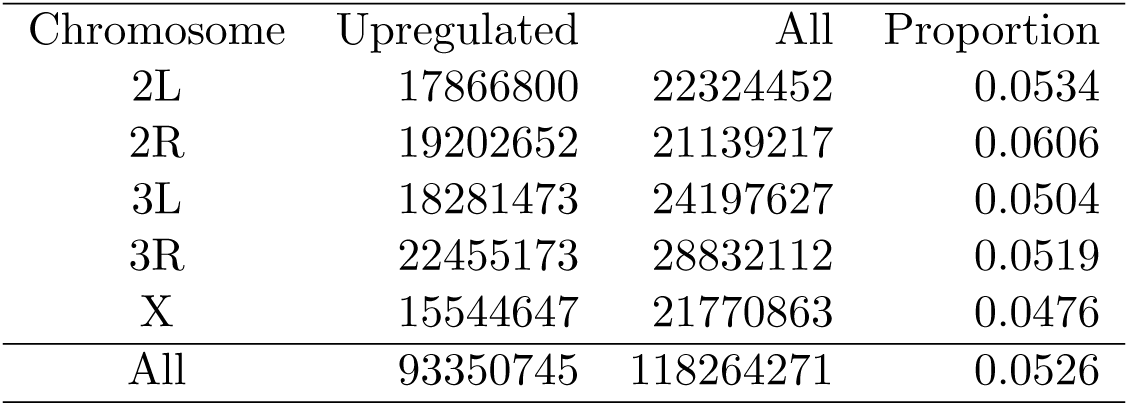
Upregulated sites genomewide

**Table S8:** Upregulated genes

| Chimeras | Tissue | $\geq 50$ bp Upregulated | Total |
| --- | --- | --- | --- |
|  | Female Carcass | 39 | 76 |
|  | Female Ovary | 40 | 76 |
|  | Male Carcass | 41 | 76 |
|  | Male Testes | 44 | 76 |
|  | All | 55 | 76 |
| Whole Gene | Tissue | $\geq 50$ bp Upregulated | Total |
|  | Female Carcass | 17 | 66 |
|  | Female Ovary | 18 | 66 |
|  | Male Carcass | 20 | 66 |
|  | Male Testes | 18 | 66 |
|  | All | 36 | 66 |

**Table S9:** Length of ‘de novo’ gene segments

| tissue | chromosome | start | stop | strain | size (bp) |
| --- | --- | --- | --- | --- | --- |
| Male Carcass | 2L | 4764717 | 4771771 | 1 | 201 |
|  | 2L | 7100699 | 7103913 | 6 | 212 |
|  | 2L | 7043543 | 7048586 | 5 | 217 |
|  | 2L | 7043543 | 7048586 | 9 | 224 |
|  | 2L | 22076307 | 22081156 | 13 | 237 |
|  | 2L | 22076307 | 22081156 | 5 | 246 |
|  | 2L | 22217615 | 22221738 | 17 | 248 |
|  | 2L | 22076307 | 22081156 | 15 | 254 |
|  | 3L | 7643207 | 7647178 | 6 | 256 |
|  | 2L | 7043543 | 7048586 | 10 | 256 |
|  | 2R | 1296122 | 1299376 | 19 | 380 |
|  | 2R | 1298866 | 1302456 | 19 | 380 |
|  | 2L | 22076307 | 22081156 | 14 | 384 |
|  | 3R | 14703209 | 14705506 | 11 | 754 |
|  | 2R | 550564 | 555698 | 13 | 1364 |
| Male Testes | 2R | 8628288 | 8637097 | 5 | 202 |
|  | 2L | 14844348 | 14850368 | 2 | 205 |
|  | 2L | 19481376 | 19484185 | 11 | 214 |
|  | 2L | 21809552 | 21814176 | 5 | 227 |
|  | 3R | 15663794 | 15666868 | 8 | 234 |
|  | 2L | 21860804 | 21864242 | 19 | 245 |
|  | X | 8626278 | 8645156 | 12 | 256 |
|  | 2R | 1296122 | 1299376 | 13 | 278 |
|  | 2R | 1298866 | 1302456 | 13 | 278 |
|  | 2L | 22076307 | 22081156 | 5 | 292 |
|  | 2R | 1296122 | 1299376 | 19 | 303 |
|  | 2R | 1298866 | 1302456 | 9 | 304 |
|  | 3R | 28773101 | 28773775 | 8 | 306 |
|  | 2L | 7043543 | 7048586 | 9 | 326 |
|  | 2R | 12531901 | 12536511 | 10 | 327 |
|  | 2L | 1858014 | 1866626 | 19 | 353 |
|  | 2L | 4764717 | 4771771 | 1 | 353 |
|  | 2R | 1298866 | 1302456 | 19 | 355 |
|  | 2L | 22229672 | 22240590 | 12 | 374 |
|  | 2L | 22076307 | 22081156 | 13 | 380 |
|  | 2R | 261487 | 266019 | 2 | 381 |
|  | 2R | 13593056 | 13597666 | 11 | 387 |
|  | 3L | 15707277 | 15731097 | 6 | 412 |
|  | 2L | 5056039 | 5058911 | 5 | 428 |
|  | 2R | 15243572 | 15249038 | 6 | 481 |
|  | 3R | 7559447 | 7567609 | 6 | 569 |
|  | 3R | 14703209 | 14705506 | 11 | 575 |
|  | 2L | 22076307 | 22081156 | 14 | 594 |
|  | 2L | 22229672 | 22240590 | 13 | 846 |
| Female Carcass | 3L | 7643207 | 7647178 | 6 | 204 |
|  | 2L | 22076307 | 22081156 | 15 | 227 |
|  | 2L | 22076307 | 22081156 | 13 | 228 |
|  | 2R | 1298866 | 1302456 | 13 | 231 |
|  | 2L | 6538411 | 6540646 | 1 | 258 |
|  | 2R | 1296122 | 1299376 | 13 | 353 |
|  | 3R | 14703209 | 14705506 | 11 | 770 |
|  | 2R | 550564 | 555698 | 13 | 1056 |
| Female Ovary | X | 21252863 | 21277771 | 14 | 343 |
|  | 2R | 1340493 | 1343865 | 6 | 686 |

**Table S10:** FPKM for recruited non-coding parental genes

| Gene | Female Carcass | Female Ovary | Male Carcass | Male Testes |
| --- | --- | --- | --- | --- |
| GE19344 | 1.11753 | 4.32018 | 0.956197 | 1.4404 |
| GE20665 | 28.1935 | 19.4317 | 29.7131 | 17.7321 |
| 2.g418 | 8.6822 | 0.0 | 3.83027 | 3.80075 |
| GE14641 | 0.00866161 | 0.0 | 3.06248 | 51.3189 |
| GE14103 | 36.4129 | 135.039 | 28.3912 | 86.1859 |
| 3.g1278 | 9.08642 | 10.7779 | 10.0527 | 11.0089 |
| GE20792 | 10.8708 | 21.321 | 8.42913 | 5.57496 |
| GE17340 | 0.77299 | 20.1854 | 3.78907 | 79.9769 |
| GE22019 | 68.3313 | 9.05024 | 89.1704 | 43.2689 |
| 2.g418 | 8.6822 | 0.0 | 3.83027 | 3.80075 |
| GE26314 | 5.25971 | 12.9278 | 3.99471 | 8.6008 |
| 4.g321 | 0.935342 | 0.405618 | 0.908753 | 0.551468 |
| 1.g396 | 0.548732 | 0.0167492 | 1.65429 | 26.874 |
| GE22133 | 14.7227 | 83.3471 | 15.9165 | 23.4822 |
| GE18873 | 7.2591 | 55.5697 | 5.87663 | 23.0839 |
| GE18174 | 58.0966 | 34.1165 | 55.9907 | 31.9375 |
| GE19410 | 13.1419 | 47.2782 | 13.6642 | 88.2691 |
| GE22569 | 0.0237261 | 0.0 | 0.0238531 | 0.419282 |
| GE15832 | 0.0205719 | 0.150601 | 0.0233789 | 0.158375 |
| 2.g1622 | 0.484331 | 0.0 | 0.083226 | 0.213026 |
| 0.g951 | 0.125324 | 0.00351728 | 0.258543 | 0.0405134 |
| GE21054 | 0.776906 | 1.85926 | 0.526995 | 1.64772 |
| 1.g5 | 2.82885 | 0.484007 | 3.67932 | 3.00005 |
| GE16826 | 8.67266 | 38.7514 | 8.17896 | 8.1714 |
| GE13040 | 0.0291575 | 0.0112303 | 0.13833 | 0.716646 |
| GE13038 | 1.44773 | 0.0 | 5.34928 | 0.369484 |
| GE21286 | 7.33772 | 35.3616 | 5.99842 | 25.6466 |
| GE12967 | 1.9533 | 5.76889 | 1.81867 | 1.42007 |
| 1.g1354 | 0.0419969 | 0.0 | 0.111257 | 0.092872 |
| GE16584 | 2.03507 | 16.8762 | 2.12453 | 5.10268 |
| GE26259 | 40.6827 | 0.128916 | 15.1555 | 10.2535 |
| GE12967 | 1.9533 | 5.76889 | 1.81867 | 1.42007 |
| GE16953 | 0.120014 | 0.537476 | 0.120596 | 0.0690305 |
| 2.g361 | 0.00773397 | 0.0 | 0.0 | 0.0192081 |
| GE26071 | 8.55841 | 0.68632 | 4.21104 | 2.41655 |
| GE16978 | 3.92049 | 17.4673 | 3.13624 | 4.64776 |
| GE13160 | 0.561831 | 0.215925 | 0.972459 | 11.6932 |
| GE15086 | 2.49297 | 0.0892037 | 7.21166 | 1.44863 |
| 2.g1732 | 0.474107 | 0.0 | 1.4353 | 0.164268 |
| GE17162 | 3.17218 | 16.647 | 3.65224 | 1.82999 |
| GE10771 | 0.0902181 | 0.011496 | 0.110188 | 0.0291091 |
| 3.g15 | 0.0 | 0.510221 | 0.0 | 0.0 |
| GE12967 | 1.9533 | 5.76889 | 1.81867 | 1.42007 |

**Table S11:** Ancestral Expression Patterns

| Tissue | Dup Mean FPKM | All Mean FPKM | Dup Median FPKM | All Median FPKM | Wilcox $W$ | $P$-value |
| --- | --- | --- | --- | --- | --- | --- |
| Ovary | 23.12815 | 16.65176 | 0.5913 | 0.3053 | 8254952 | $3.291 \times 10^{-4}$ |
| Female Carcass | 19.0621 | 16.8729 | 2.6573 | 1.3851 | 8884288 | $2.282 \times 10^{-16}$ |
| Testes | 17.78303 | 15.1603 | 3.3762 | 1.9954 | 8743698 | $7.368 \times 10^{-13}$ |
| Male Carcass | 20.34798 | 17.2835 | 9040304 | 3.3519 | 1.9687 | $2.2 \times 10^{-16}$ |

**Table S12:** Ancestral Expression Patterns for Variants in ≤ 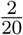 Strains

| Tissue | Dup Mean FPKM | All Mean FPKM | Wilcox $W$ | $P$-value |
| --- | --- | --- | --- | --- |
| Ovary | 24.170 | 16.650 | 7207434 | $4.148 \times 10^{-5}$ |
| Female Carcass | 18.910 | 16.879 | 7686539 | $2.694 \times 10^{-15}$ |
| Testes | 17.300 | 15.160 | 7588718 | $1.045 \times 10^{-12}$ |
| Male Carcass | 20.912 | 17.284 | 7844719 | $2.2 \times 10^{-16}$ |

**Table S13:** Sample strains surveyed

| Stock Number | Strain |
| --- | --- |
| 14021-0261.01 | Reference |
| 14021-0261.39 | CY04B |
| 14021-0261.40 | CY08A |
| 14021-0261.41 | CY17C |
| 14021-0261.42 | CY20A |
| 14021-0261.43 | CY21B3 |
| 14021-0261.44 | CY22B |
| 14021-0261.47 | NY48 |
| 14021-0261.48 | NY56 |
| 14021-0261.49 | NY62 |
| 14021-0261.50 | NY65 |
| 14021-0261.51 | NY66-2 |
| 14021-0261.52 | NY73 |
| 14021-0261.53 | NY81 |
| 14021-0261.54 | NY85 |
| N/A | CY28A |

**Table S14:** Upregulated genes using Baum-Welch transition probabilities

| Chimeras | Tissue | Upregulated | Total |
| --- | --- | --- | --- |
|  | Female Carcass | 5 | 76 |
|  | Female Ovary | 10 | 76 |
|  | Male Carcass | 10 | 76 |
|  | Male Testes | 9 | 76 |
|  | All | 22 | 76 |
| Whole Gene | Tissue | Upregulated | Total |
|  | Female Carcass | 3 | 66 |
|  | Female Ovary | 2 | 66 |
|  | Male Carcass | 1 | 66 |
|  | Male Testes | 1 | 66 |
|  | All | 5 | 66 |

**Figure S1:**
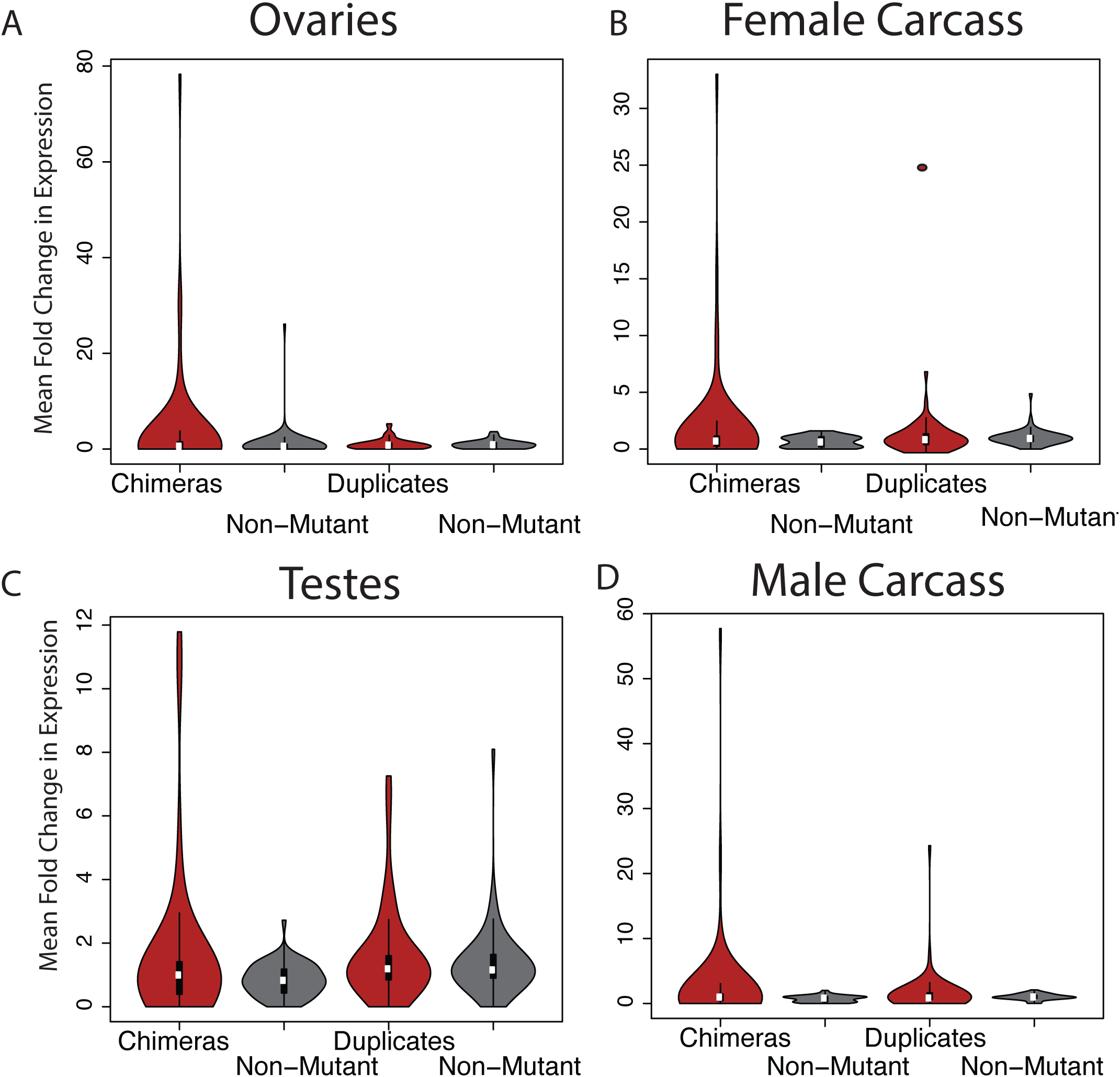
Mean fold change for chimeric genes in sample strains vs. reference for strains containing chimeras or whole gene duplicates (red) and unmutated sample strains for the same regions (grey). Chimeric genes are more likely to result in high mean fold change than unmutated counterparts in all tissues. Whole gene duplicates create multifold expression changes more rarely.

**Figure S2:**
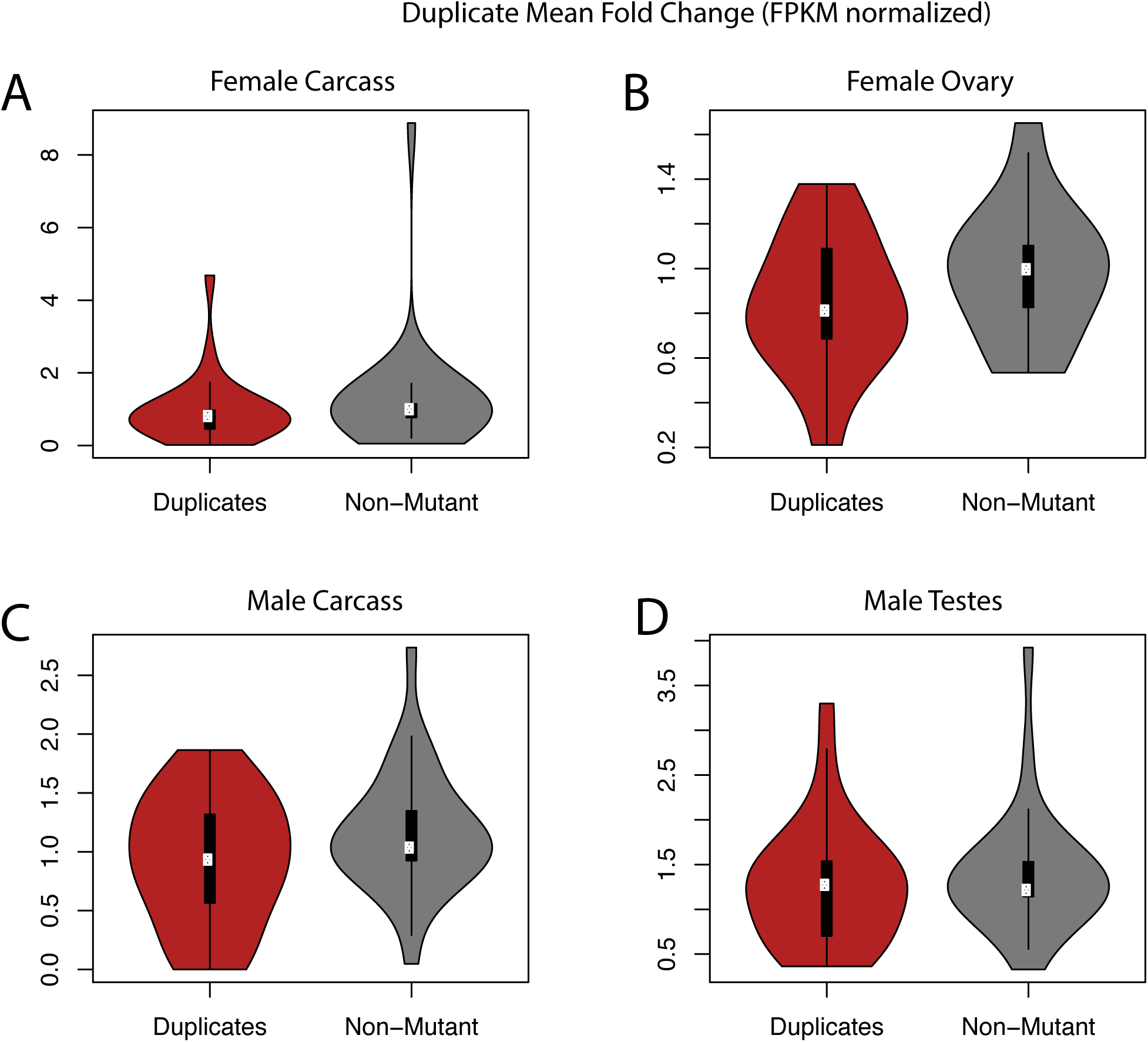
Mean fold change using FPKM normalized data for chimeric genes in sample strains vs. reference for strains containing chimeras or whole gene duplicates (red) and unmutated sample strains for the same regions (grey). Chimeric genes are more likely to result in high mean fold change than unmutated counterparts in all tissues. Whole gene duplicates create multifold expression changes more rarely.

**Figure S3:**
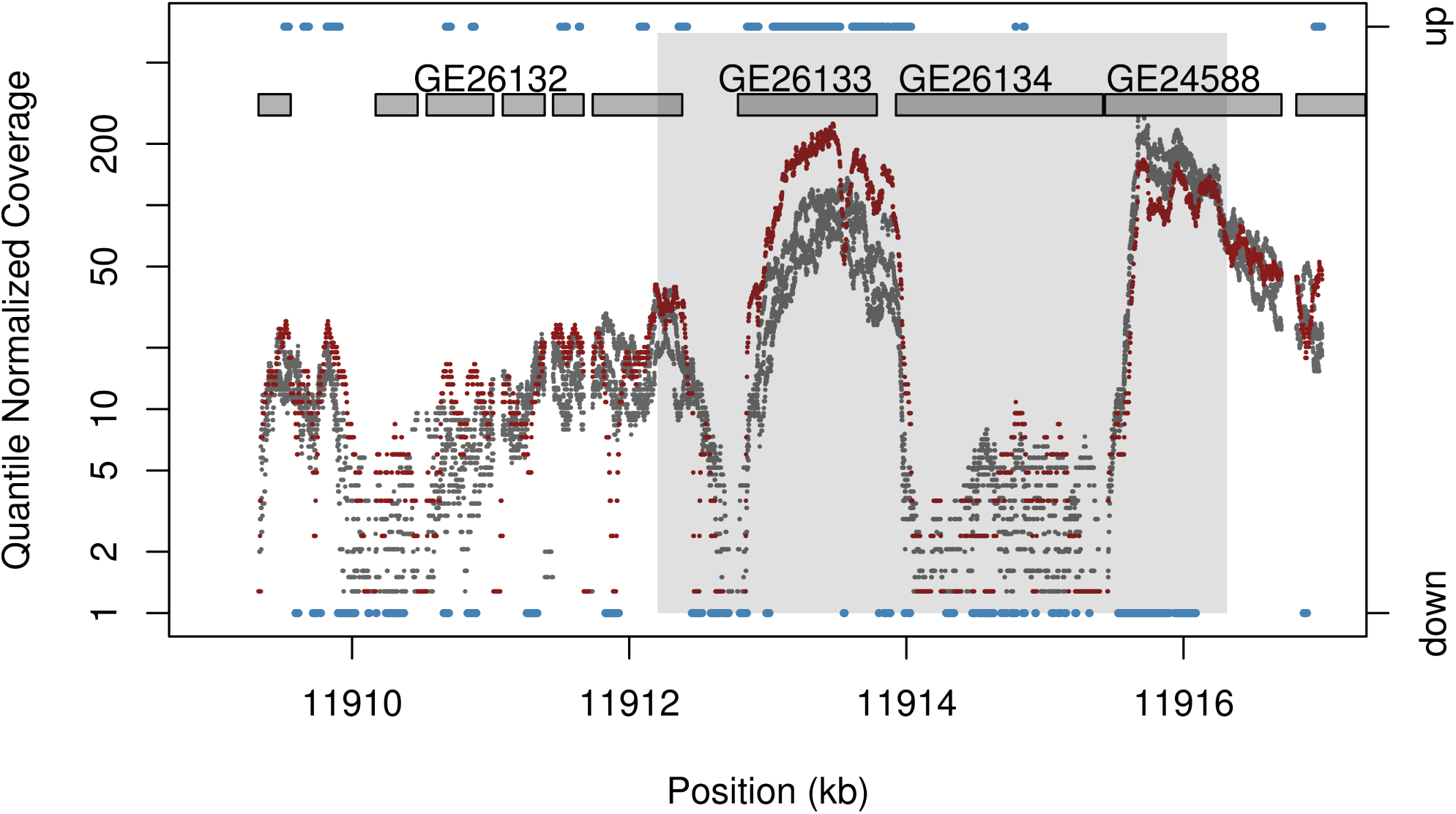
Expression change in a sample strain containing a whole gene duplication of *GE26133* (reference FPKM=22.0725, sample FPKM=58.6217, uncorrected *P* = 0.00263417, corrected *P* = 0.0420917). The tandem duplication also captures the entire gene sequence of *GE26134*, as well as portions of *GE26132* and *GE24588*. The duplicate exhibits greater than two-fold expression of *GE26133* in the sample strain containing the duplication. It is unclear whether the expression change is a direct consequence of duplication, secondary mutation, environmental effects, or other stochastic variation in expression.

**Figure S4:**
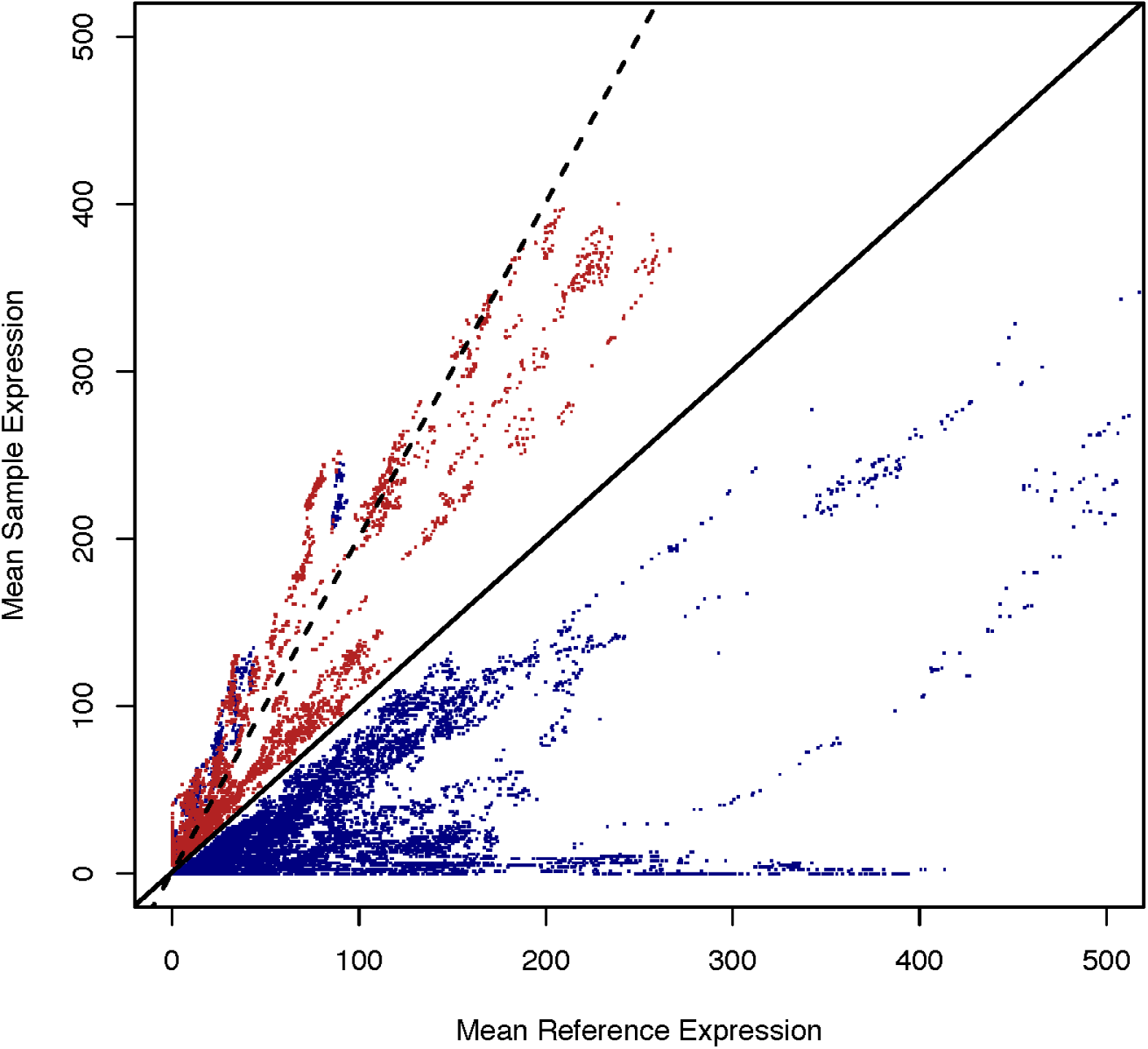
HMM Performance in quantile normalized coverage data. Quantile normalized coverage in a single sample vs. the mean of quantile normalized coverage in the reference for sites with upregulated sequence are plotted in red, while that of down regulated sequence is shown in blue for 500,000 bp beginning at 6.5 Mb on chromosome 3L for sites with quantile normalized coverage ≤ 500. Sites with no expression change identified using the HMM are not shown. The case of equal expression is shown with the black solid line, while two-fold coverage increase in the sample are indicated with the dashed line. Even modest increases in expression can be identified with the HMM, suggesting that its ability to detect site level differences in high coverage RNA-seq data is high.

**Figure S5:**
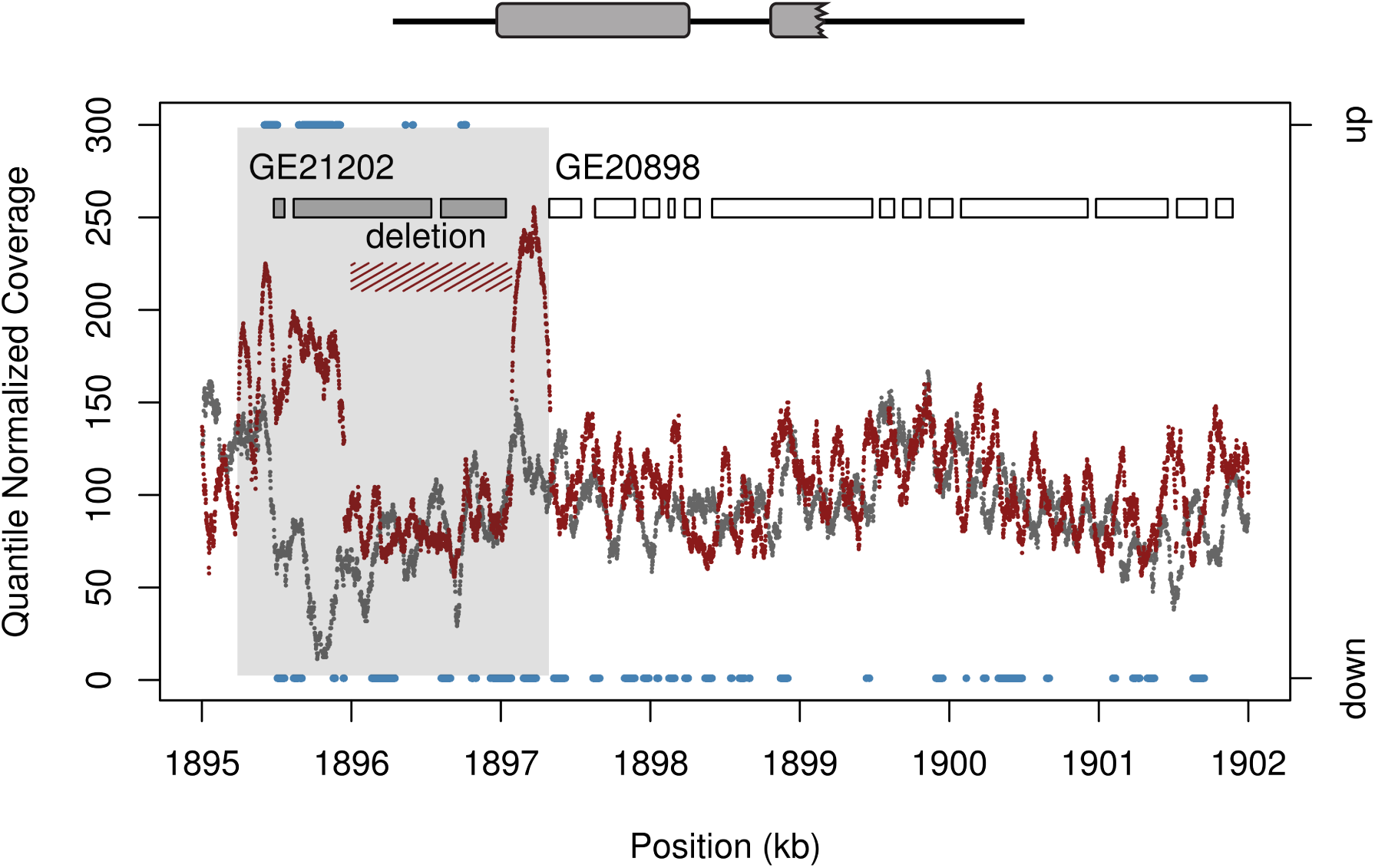
Genomic DNA sequencing coverage in the sample (red) and resequenced reference (grey) (14) and RNA-seq HMM Expression output for a region experiencing a secondary deletion after duplication. The deleted segment is supported by a decrease in genome coverage as well as 104 long-spanning Illumina sequencing reads. Coverage increases two-fold to three-fold in the duplicated segment, and is not supportive of higher level copy number that might explain the increase in expression as defined by RNA-seq data. HMM output for the region with increased expression in RNA-seq data is shown in blue, for comparison. The region the gene segment with the expression change corresponds well with the region displaying elevated genomic sequencing coverage given the structure of ancestral gene models (see Figure 3).

**Figure S6:**
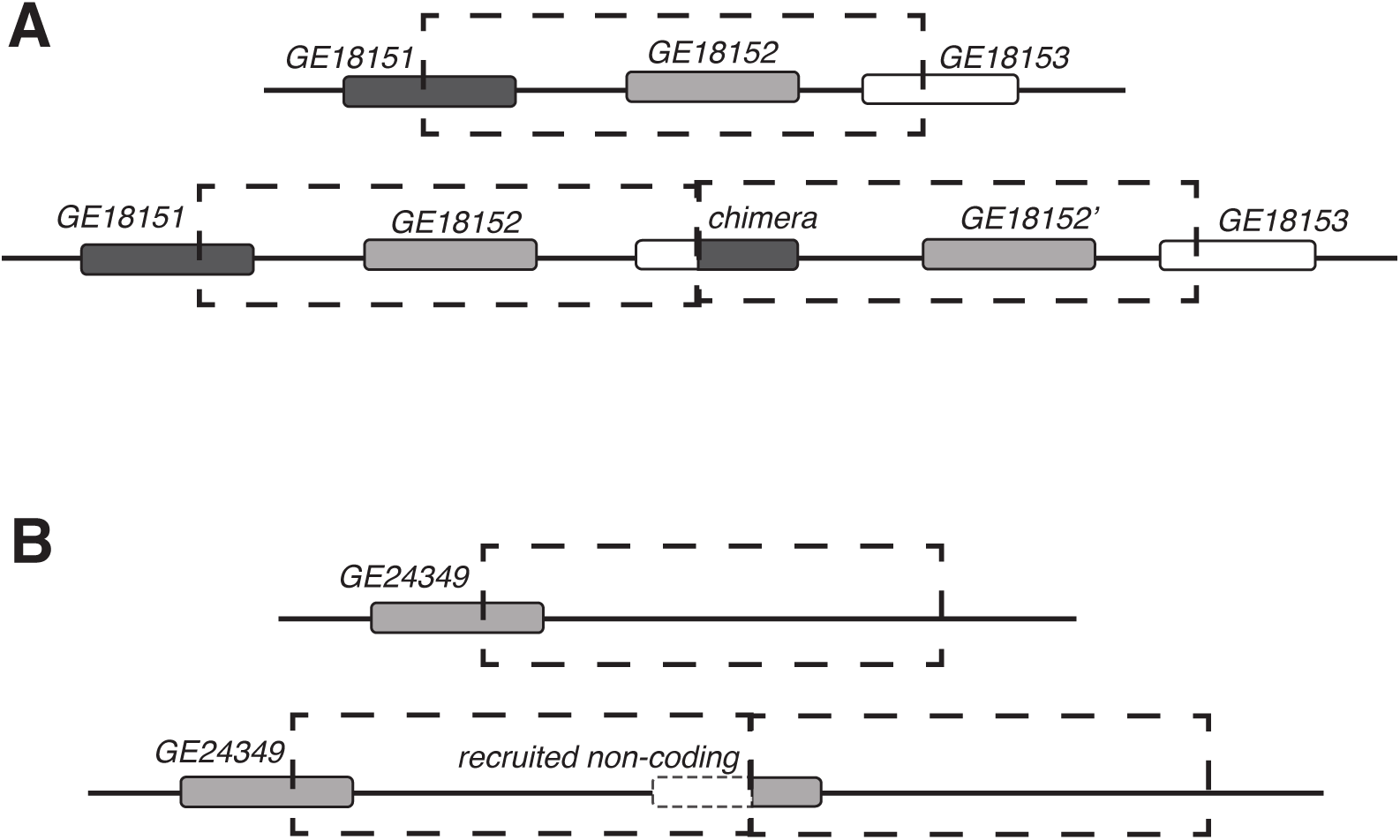
Formation of alternative gene structures through tandem duplications. A) A tandem duplication captures the 5ʹ segment of *GE18453* and the 3ʹ segment of *GE18451*. The tandem duplication unites these gene segments to form a novel open reading frame distinct from the parental genes. Shuffling of regulatory elements in the 5ʹ and 3ʹ ends results in a new regulatory profile for the chimera. The tandem duplication also copies the full gene sequence of *GE18452*. B) A tandem duplication captures the 5ʹ end of *GE24349*, placing it next to previously untranscribed sequence. The promoter and UTR of *GE24349* drives expression of a previously untranscribed region.

**Figure S7:**
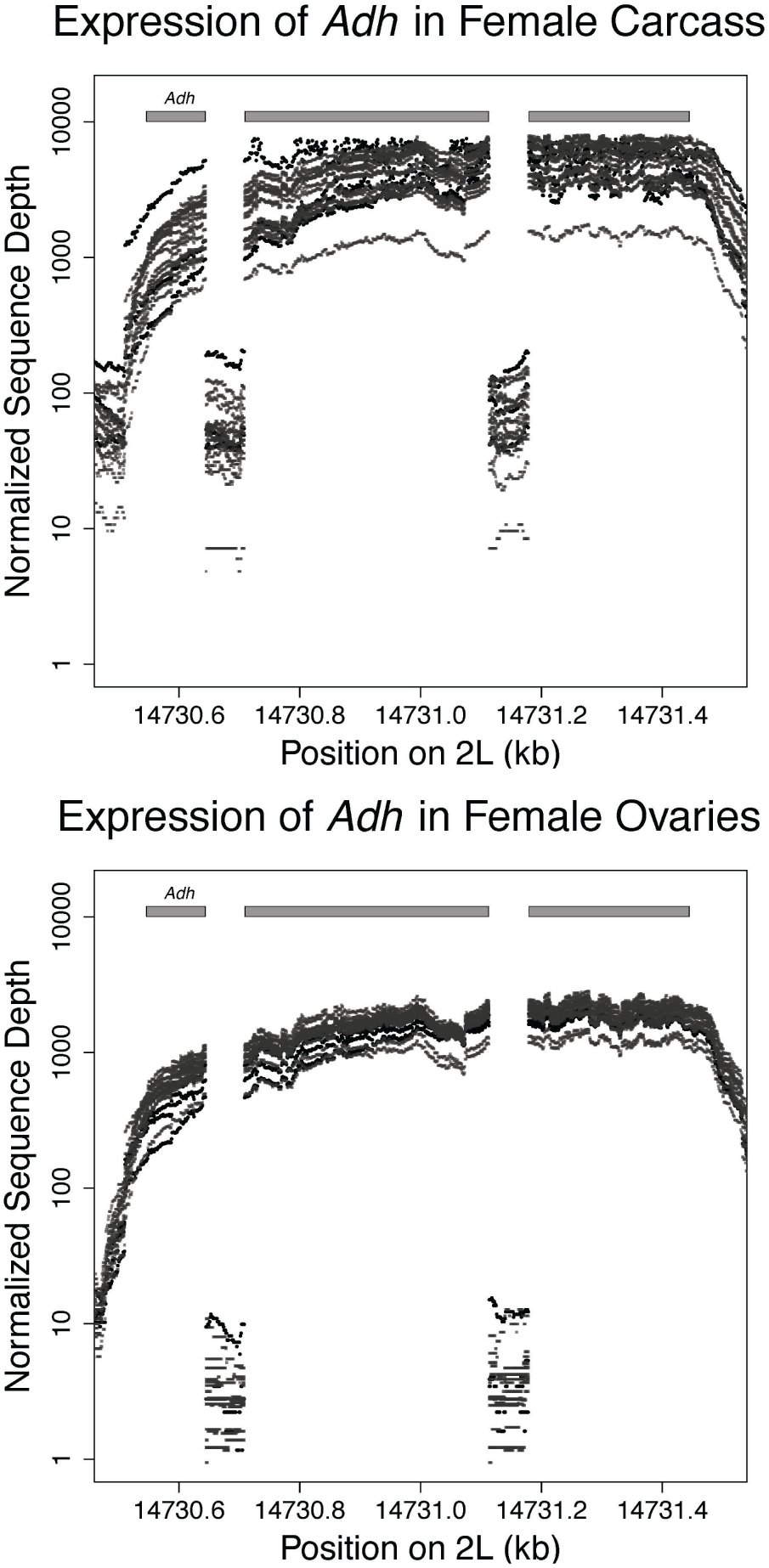
Normalized coverage in RNA-seq Data for *Adh* in 15 sample strains and 3 replicates of the reference. RNA-seq data shows differentiation between intron and exon sequence and spans the entire length of the the transcript.

**Figure S8:**
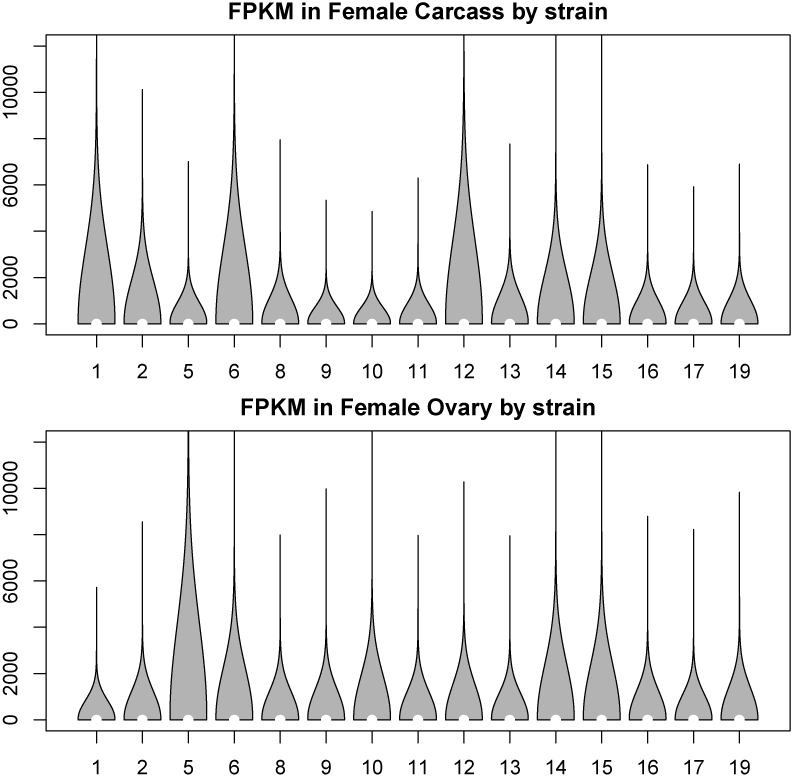
FPKM values in RNA-seq data in female tissues for 15 sample strains. Coverage varies across strains, but is generally high with thousands of reads for the most highly expressed genes. To reduce variability in coverage and generate more robust differential expression calls, we quantile normalized coverage inputs for the HMM.

**Figure S9:**
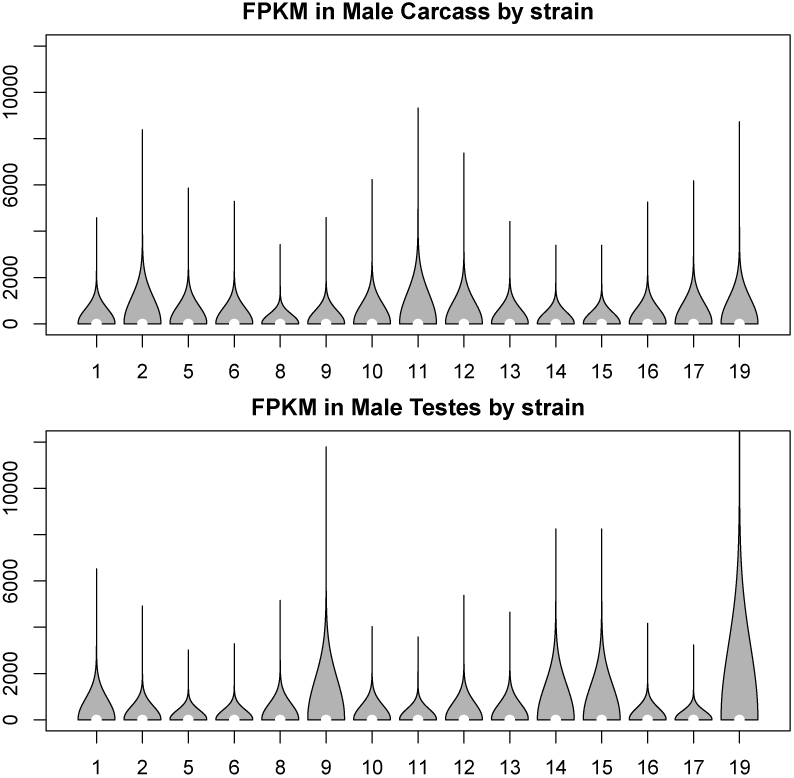
FPKM values in RNA-seq data in male tissues for 15 sample strains. Coverage varies across strains, but is generally high with thousands of reads for the most highly expressed genes. To reduce variability in coverage and generate more robust differential expression calls, we quantile normalized coverage inputs for the HMM.

